# Hydration-dependent phase separation of a prion-like protein regulates seed germination during water stress

**DOI:** 10.1101/2020.08.07.242172

**Authors:** Yanniv Dorone, Steven Boeynaems, Benjamin Jin, Flavia Bossi, Eduardo Flores, Elena Lazarus, Emiel Michiels, Mathias De Decker, Pieter Baatsen, Alex S. Holehouse, Shahar Sukenik, Aaron D. Gitler, Seung Y. Rhee

## Abstract

Many organisms evolved strategies to survive and thrive under extreme desiccation. Plant seeds protect dehydrated embryos from a variety of stressors and can even lay dormant for millennia. While hydration is the key trigger that reactivates metabolism and kick-starts germination, the exact mechanism by which the embryo senses water remains unresolved. We identified an uncharacterized *Arabidopsis thaliana* prion-like protein we named FLOE1, which phase separates upon hydration and allows the embryo to sense water stress. We demonstrate that the biophysical states of FLOE1 condensates modulate its biological function *in vivo* in suppressing seed germination under unfavorable environments. We also find intragenic, intraspecific, and interspecific natural variations in phase separation propensity of FLOE1 homologs. These findings demonstrate a physiological role of phase separation in a multicellular organism and have direct implications for plant ecology and agriculture, especially the design of drought resistant crops, in the face of climate change.

## MAIN TEXT

Plant seeds are specialized propagation vectors that can mature to a quiescent, desiccated state, allowing them to remain viable in harsh conditions anywhere from a few years to millennia (*1, 2*). Although water is essential for life, plant embryos can survive extreme desiccation by accumulating protective molecules that profoundly change their cellular biophysical properties (*3, 4*). Upon the uptake of water, called imbibition, seeds rapidly undergo a cascade of biochemical and mechanical events and the resumption of cellular activities (*5, 6*). Seeds can endure multiple hydration-dehydration cycles while remaining viable and desiccation tolerant (*7*). However, once committed to germination, they can no longer revert to their stress tolerant state (*5*). Thus, poor timing of germination can severely limit seedling survival (*8*), especially in times of drought. Despite the fundamental importance of germination control for plant biology and agriculture, the molecular details that underpin this decision remain incompletely understood.

Given limited water availability dramatically alters protein solubility and seeds are known to undergo a cytoplasmic liquid-to-glass transition during maturation (*3, 4*), we investigated how seed proteins might have adapted to these extreme conditions. We re-analyzed existing *Arabidopsis thaliana* transcriptomics data and found 449 protein-coding genes that are expressed at higher relative levels in dry seeds compared to other tissues (Fig. 1A, Table S1) (*9, 10*). These seed-enriched proteins had a different amino acid compositional profile (Fig. 1B) and were enriched for regions of structural disorder compared to the rest of the proteome (Fig. 1C). Intrinsically disordered proteins (IDPs) have emerged as key players in cell biology that, among many roles, orchestrate how cells organize themselves and their contingent biochemical reactions into discrete membraneless compartments by a process called liquid-liquid phase separation (LLPS) (*11, 12*). A subset of IDPs harbor a class of domains known as prion-like domains (PrLD), and we identified 14 proteins with PrLDs in the seed-enriched proteins (Fig. 1D). PrLDs were first identified in fungal prions and more recently were shown to drive reversible protein phase separation in diverse eukaryotic species (*13*). In yeast, PrLDs give rise to phenotypic diversity via a proposed bet-hedging strategy to help cope with and survive in a fluctuating environment (*14*). All but one of these PrLD-containing seed-enriched proteins had annotated functions or domains related to nucleic acid metabolism. The one that did not, AT4G28300, was an uncharacterized plant-specific protein, which we named FLOE1 (inspired by the second movement of *Glassworks* by Philip Glass as well as the definition of floe being ‘a sheet of floating ice’, which is a phase-separated body of water).

**Figure 1:**
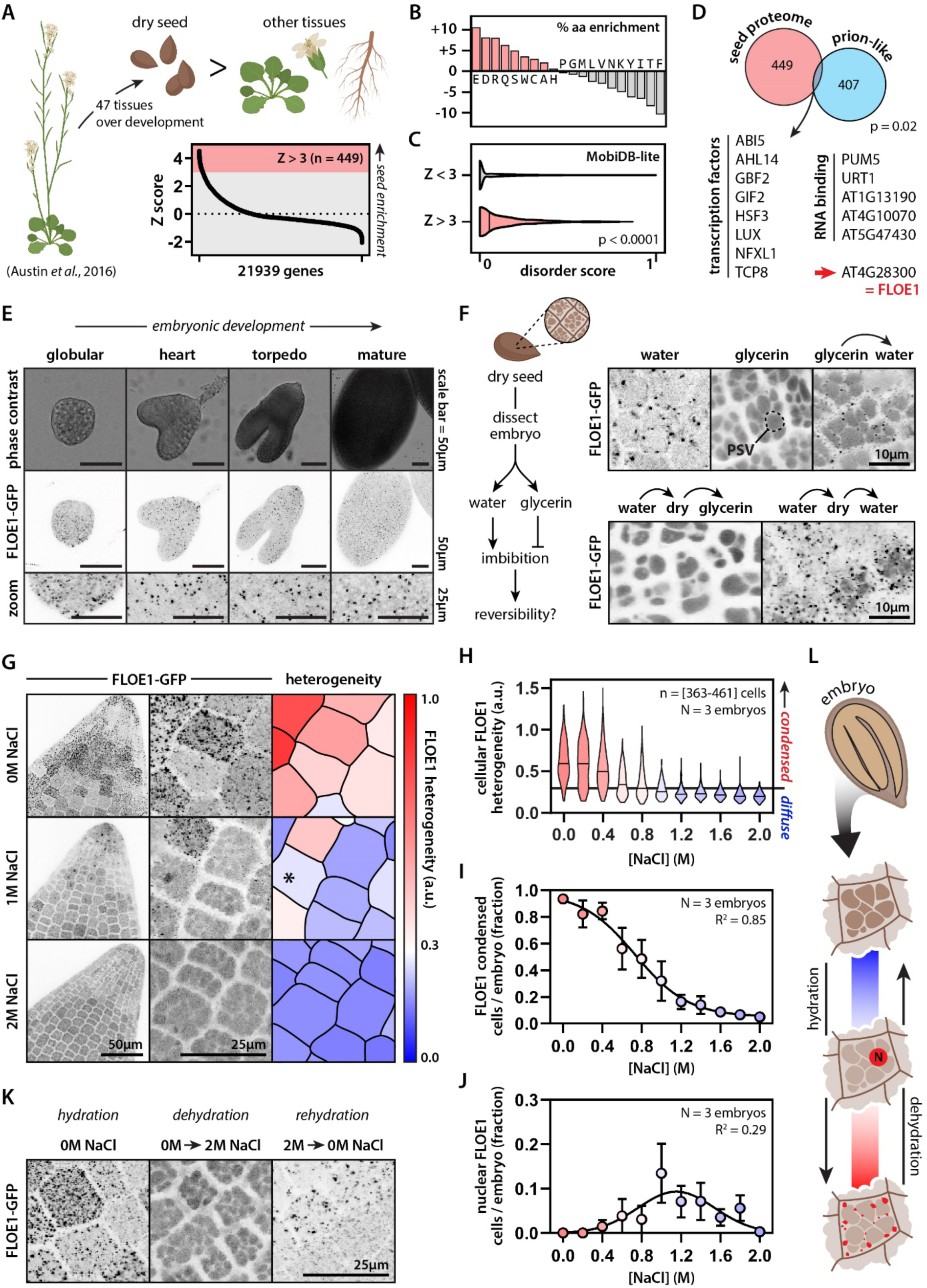
FLOE1 is an uncharacterized seed protein that undergoes biomolecular condensation in a hydration-dependent manner. (A) Identification of genes enriched in dry *Arabidopsis* seeds. (B-C) The seed proteome is enriched for specific amino acids (B) and intrinsic disorder (C). Mann-Whitney. (D) The seed proteome is enriched for prion-like proteins. Binomial test. AT4G28300 is an uncharacterized prion-like protein, which we name here FLOE1. (E) FLOE1p:FLOE1-GFP is expressed during embryonic development and forms condensates. (F) FLOE1-GFP forms condensates in embryos dissected from dry seed in a hydration-dependent and reversible manner. Embryonic cotyledons are shown. PSV denotes autofluorescent protein storage vacuoles that are more prominent in the dry state than in the hydrated state (see also Fig. S2C). (G) Cell-to-cell variation in subcellular FLOE1-GFP heterogeneity in response to salt. Radicles are shown. * denotes nuclear localization. (H) Quantification of cellular FLOE1 heterogeneity as a function of salt concentration. Black line denotes the 95^th^ percentile of the 2M NaCl heterogeneity distribution. (I) Quantification of the percentage of cells per radicle that show FLOE1 condensation as a function of salt concentration. Mean ± SEM. Four-parameter dose-response fit. (J) Quantification of the percentage of cells per radicle that show FLOE1 nuclear localization as a function of salt concentration. Mean ± SEM. Gaussian fit. (K) FLOE1-GFP condensation exhibits reversibility between high and no salt treatment. Radicles are shown. (L) Scheme highlighting different FLOE1 behaviors upon imbibition.

FLOE1 accumulates during embryo development and its expression peaks in the mature desiccated state (Fig. S1). We generated transgenic *A. thaliana* lines expressing *FLOE1-GFP* under control of its endogenous promoter and with its non-coding sequences intact. FLOE1 formed cytoplasmic condensates during embryonic development (Fig. 1E, Fig. S2A) and in embryos dissected from dry seeds (Fig. 1F, Fig. S2B). However, when we dissected dry seeds in glycerin instead of water (to mimic the desiccated environment), FLOE1 did not form condensates and was localized diffusely (Fig. 1F, Fig. S2C-D). When we transferred these embryos from glycerin to water, FLOE1 condensates spontaneously appeared (Fig. 1F) and were fully reversible with repeated hydration-dehydration cycles (Fig. 1F). Pre-treating seeds with the translation inhibitor cycloheximide did not affect the formation of FLOE1 condensates, indicating that they are distinct from stress granules and processing bodies (*15*), and that their emergence did not depend on FLOE1 translation upon imbibition (Fig. S2E). To directly test whether FLOE1 forms condensates in response to changes in water potential, we dissected embryos in solutions of varying concentrations of salt, mannitol, or sorbitol (Fig. 1G-I; Fig. S2F-G). High concentrations of salt resembled dry conditions in that embryos lacked visible FLOE1 condensates (Fig. 1G-I). Lowering the salt concentration resulted in a gradual emergence of condensates, which was highly variable at the cell-to-cell (Fig. 1G-H) and tissue levels (Fig. 1I). In intermediate salt concentrations, we observed a small number of cells with apparent nuclear localization of FLOE1 (Fig. 1J), suggesting this could be a behavior associated with early steps of imbibition, before the majority of the protein condenses in the cytoplasm. Similar to our observations with repeated hydration-dehydration cycles, FLOE1 condensation was also reversible by moving the embryos back and forth between solutions of high or no salt (Fig. 1K). We conclude that FLOE1 forms cytoplasmic condensates in response to changes in water potential (Fig. 1L).

Numerous yeast proteins undergo oligomerization or phase separation upon stress-induced quiescence (*16*) but to our knowledge FLOE1 is the first example of a protein undergoing biomolecular condensation upon *release* from the quiescent state. To define the mechanism by which FLOE1 undergoes this switch, we dissected the molecular grammar underlying this behavior. We first expressed and purified MBP-tagged FLOE1 protein from *Escherichia coli* to determine if it undergoes phase separation *in vitro*. Upon cleavage of the MBP tag with TEV protease, FLOE1 spontaneously demixed to form droplets (Fig. 2A, Fig. S3A-B) that enriched FLOE1 but excluded MBP (Fig. S3C), similar to the behavior of other phase-separating proteins (*17*). FLOE1 harbors a predicted coiled-coil domain and a conserved plant-specific domain of unknown function (DUF1421) (Fig. 2B). Disorder prediction algorithms identified another predicted folded region and two different disordered regions, one enriched for amino acids aspartic acid and serine (DS-rich) and the other enriched for glutamine, proline, and serine (QPS-rich). We heterologously expressed FLOE1 in two orthogonal systems, tobacco leaves (Fig. 2C, Fig. S4) and the human osteosarcoma cell line U2OS (Fig. 2D). In these two systems, as already shown in *A. thaliana*, FLOE1 formed spherical condensates, providing independent platforms for interrogating the molecular drivers of condensation. We systematically deleted each domain of FLOE1 and assayed the impact on cytoplasmic condensation (Fig. 2C-E). In both tobacco and human cells, mutants lacking either the short coiled-coil domain or DUF1421 behaved identically to the wildtype protein (Fig. 2C-E). Deletion of the other domains altered FLOE1 condensation (Fig. 2C-E). Deletion of the predicted folded domain, which we refer to as the nucleation domain, abolished cytoplasmic condensation, resulting in a fraction of the protein redistributing to the nucleus. Folded oligomerization domains play important roles in nucleating phase separation of several IDPs (*12*). Indeed, expression of chimeric fusion proteins revealed that this domain is sufficient to nucleate phase separation of different PrLDs (Fig. 2F). All-atom simulations and homology-based modeling suggest that the nucleation domain adopts a trimeric coiled coil conformation (Fig. S5), providing the necessary multivalent interactions required for nucleating protein phase separation.

**Figure 2:**
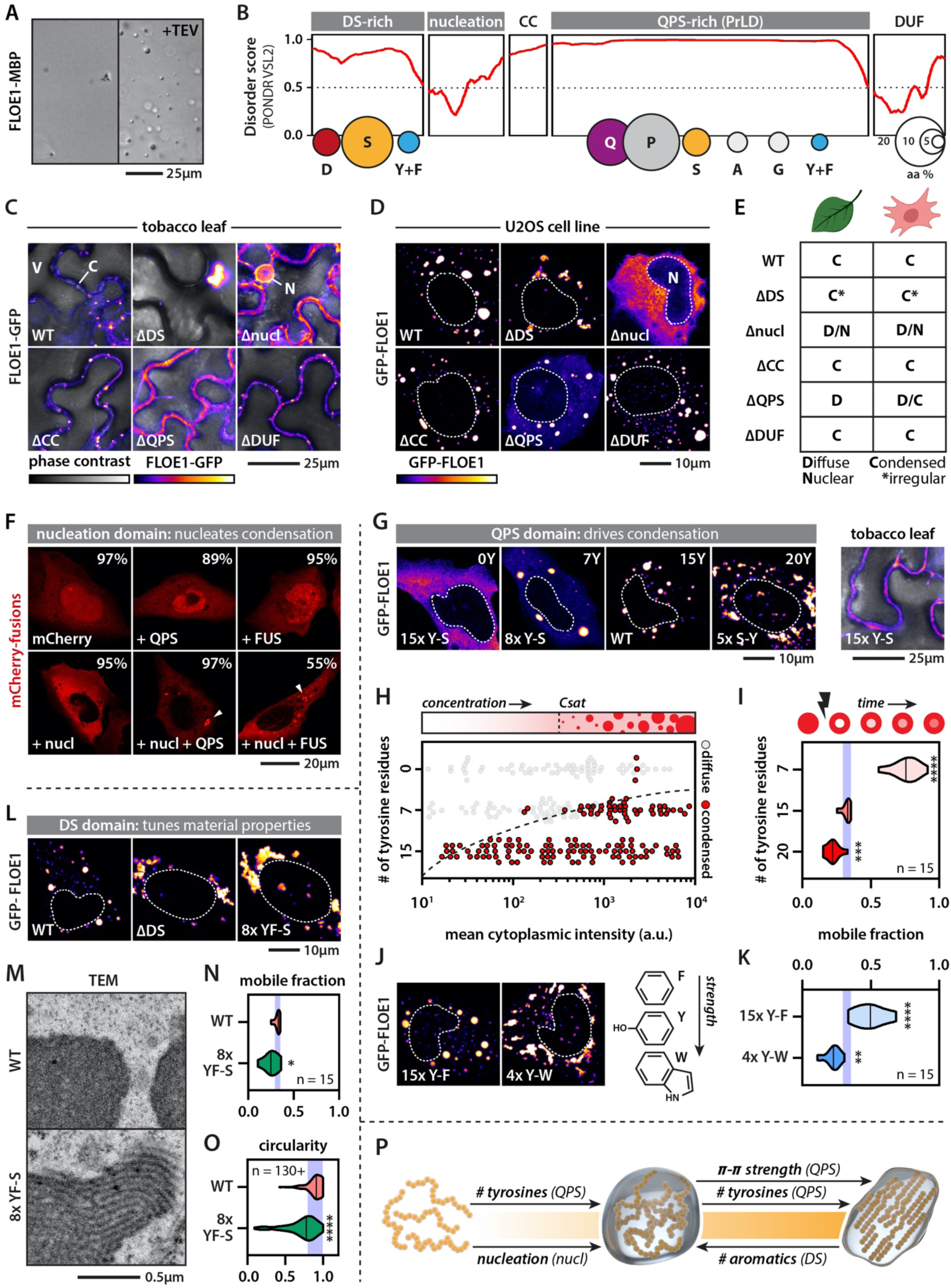
Molecular dissection of FLOE1 phase separation. (A) Recombinant MBP-FLOE1 phase separates in the test tube upon MBP cleavage with TEV protease. Irregular small aggregates can be seen pre-cleavage highlighting FLOE1 aggregation-propensity. (B) FLOE1 domain structure. CC = predicted coiled coil, DUF = DUF1421. Balloon plots show amino acid composition of the disordered domains. (C-D) Expression of FLOE1 domain deletion mutants in tobacco leaves (C) and human U2OS cells (D). V = vacuole, C = cytoplasm, N = nucleus. (E) Summary of FLOE1 behavior in tobacco leaves and human cells. (F) Chimeric proteins containing both the FLOE1 nucleation domain and the PrLDs from FLOE1 (QPS) or the human FUS protein form cytoplasmic condensates. Percentages display number of cells lacking or containing condensates. Average of three experiments. Arrowheads point at cytoplasmic condensates. (G) The number of QPS tyrosine residues alters FLOE1 phase separation in human cells and tobacco leaves. (H) FLOE1 phase diagram as a function of concentration and number of QPS tyrosines. (I) Number of QPS tyrosines affects intracondensate FLOE1 dynamics. Mobile fraction as assayed by FRAP is shown. One-way ANOVA. (J-K) QPS tyrosine-phenylalanine and tyrosine-tryptophan substitutions alter condensate morphology (J) and intracondensate dynamics compared to WT (K). One-way ANOVA. (L) DS deletion or DS tyrosine/phenylalanine-serine substitutions alter condensate morphology. (M) TEM shows that mutant DS FLOE1 condensates have filamentous substructure that is absent in the WT. U2OS cells. (N) DS tyrosine/phenylalanine-serine substitutions alter intracondensate dynamics. Student’s t-test. (O) DS tyrosine/phenylalanine-serine substitutions alter condensate morphology. Mann-Whitney. (P) Scheme summarizing synergistic and opposing roles of FLOE1 domains on the material property spectrum. * p-value < 0.05, ** p-value < 0.01, *** p-value < 0.001, **** p-value < 0.0001. Purple band denotes WT mean ± SD (I, K, N, O)

In line with their role in driving phase separation of other prion-like proteins, deletion of the QPS PrLD reduced condensate formation (Fig. 2C-E). Consistent with the emerging sticker-spacer framework for PrLDs (*18, 19*), the QPS PrLD has regularly spaced aromatic tyrosine residues along its sequence that may act as attractive stickers (Fig. S6). Substituting tyrosine residues for serines (Y-S) decreased condensate formation in both human and tobacco cells in a dose-dependent manner (Fig. 2G, Fig. S6). By mapping out a phase diagram (Fig. 2H) and probing the molecular dynamics using fluorescence recovery after photobleaching (Fig. 2I) of Y-S and S-Y mutants, we confirmed that the number of tyrosines determines both the saturation and gelation concentration of FLOE1 condensates, consistent with what has been shown for other PrLDs (*19*). These findings provide further evidence that FLOE1 condensates form via LLPS, and increasing its multivalency drives gelation into more solid-like irregular assemblies. While changing the number of stickers can drive a liquid-to-gel transition, altering sticker strength may also alter the gelation concentration. Substituting tyrosines for weaker (phenylalanine) or stronger (tryptophan) aromatic residues affected both condensate morphology and intracondensate FLOE1 dynamics in a predictable manner (Fig. 2J-K, Fig. S6). While increasing the stickiness of the QPS PrLD induced gelation of FLOE1, this was also the case for deletion of the N-terminal DS domain (Fig. 2C-E, 2L). Surprisingly, serine substitution of aromatic residues in the DS domain had a similar effect to DS deletion (Fig. 2L), which suggests that the aromatic residues in the DS and QPS domains have opposing functions. Intriguingly, electron microscopy revealed that the gel-like condensates formed via DS-8xFY-S had a regular striated substructure, and were strikingly distinct from the homogeneous wildtype condensates (Fig. 2M, Fig S7), consistent with their reduced dynamics and morphological change (Fig. 2N-O). This striated substructure is reminiscent of synaptonemal complexes that have also been proposed to form through phase separation of a coiled coil protein (*20*). Thus, synergistic and opposing molecular forces tightly regulate FLOE1’s biophysical phase behavior, and changing this balance allows its properties to be experimentally toggled between dilute, liquid droplet and solid gel states (Fig. 2P).

We next asked whether these various physical states of FLOE1 have a functional role in germination. Lines carrying the knockout allele *floe1-1* did not show any obvious developmental defects, and *floe1-1* seeds had the same size, shape and weight as the wildtype (Fig. S8A). *floe1-1* seeds germinated indistinguishably from wildtype seeds under standard conditions (Fig. S8B) but had higher germination rates under conditions of water deprivation induced by salt (Fig 3A, Fig. S8C) or mannitol (Fig. S8C). We confirmed that these phenotypes were caused by mutations in *FLOE1* using independent lines carrying CRISPR-Cas9 *FLOE1* deletion alleles and *floe1-1* lines complemented with the wildtype allele (Fig. S8C-F). Germination during stressful environmental conditions is risky for a plant and can reduce fitness. Indeed, seedlings displayed developmental defects or eventually died under these conditions (Fig. 3B, Fig. S8G), whereas ungerminated seeds retained full germination potential upon stress alleviation (Fig. 3C), in line with bet-hedging strategies in stressed seeds (*21–23*). To probe the role of FLOE1 condensates in stress-mediated attenuation of germination, we first asked whether the condensates would reform upon alleviation from salt stress. Whereas ungerminated salt-stressed seeds were largely devoid of FLOE1 condensates, even after 15 days of incubation, alleviating salt stress induced their robust appearance (Fig. 3D, Fig. S8H). This shows that FLOE1 phase separates during physiologically relevant conditions *in vivo*. To directly test whether FLOE1’s function in seed germination depends on its ability to undergo phase separation, we generated *A. thaliana* lines carrying wildtype or different FLOE1 domain deletion mutants in the *floe1-1* null background (Fig. 3E-F). These mutants behaved identically in *A. thaliana* embryos as they did in human and tobacco cells (Fig. 2C-E). The ΔQPS mutant was unable to phase separate upon imbibition (Fig. 3F), whereas the ΔDUF mutant formed condensates similar to the wildtype (Fig. 3F-H). In contrast, the ΔDS mutant formed condensates that were much larger than those formed by the wildtype (Fig. 3F-H), consistent with what we observed in tobacco and human cells. The ΔDS mutant also seemed to have lost some of its hydration-dependency (Fig. 3I), consistent with its solid-like biophysical properties. We assayed germination rates under salt stress and found that removing the QPS domain resulted in FLOE1 loss of function (Fig. 3J, Fig. S9A-C). Surprisingly, removing the DS domain resulted in a greatly enhanced germination rate under stress, surpassing even that of the *floe1-1* null mutant, indicating that ΔDS likely functions as a gain-of-function mutation (Fig. 3J, Fig. S9A-C). Even under standard conditions, the ΔDS mutant displayed faster germination rates (Fig. S9D). In the evolutionary game theory framework (*21–23*), this ΔDS mutant behaves like a “high-stakes gambler”—it perceives the risk of germination under stress (e.g. seedling dying) to be lower than the chance of there being a change in the environment (e.g. increased rainfall). Thus, FLOE1 seems to function as a water stress-dependent “resistor” in the signaling cascade that triggers the initiation of germination upon imbibition, tuning bet-hedging strategies at this crucial step of a seed’s life (Fig. 3K).

**Figure 3:**
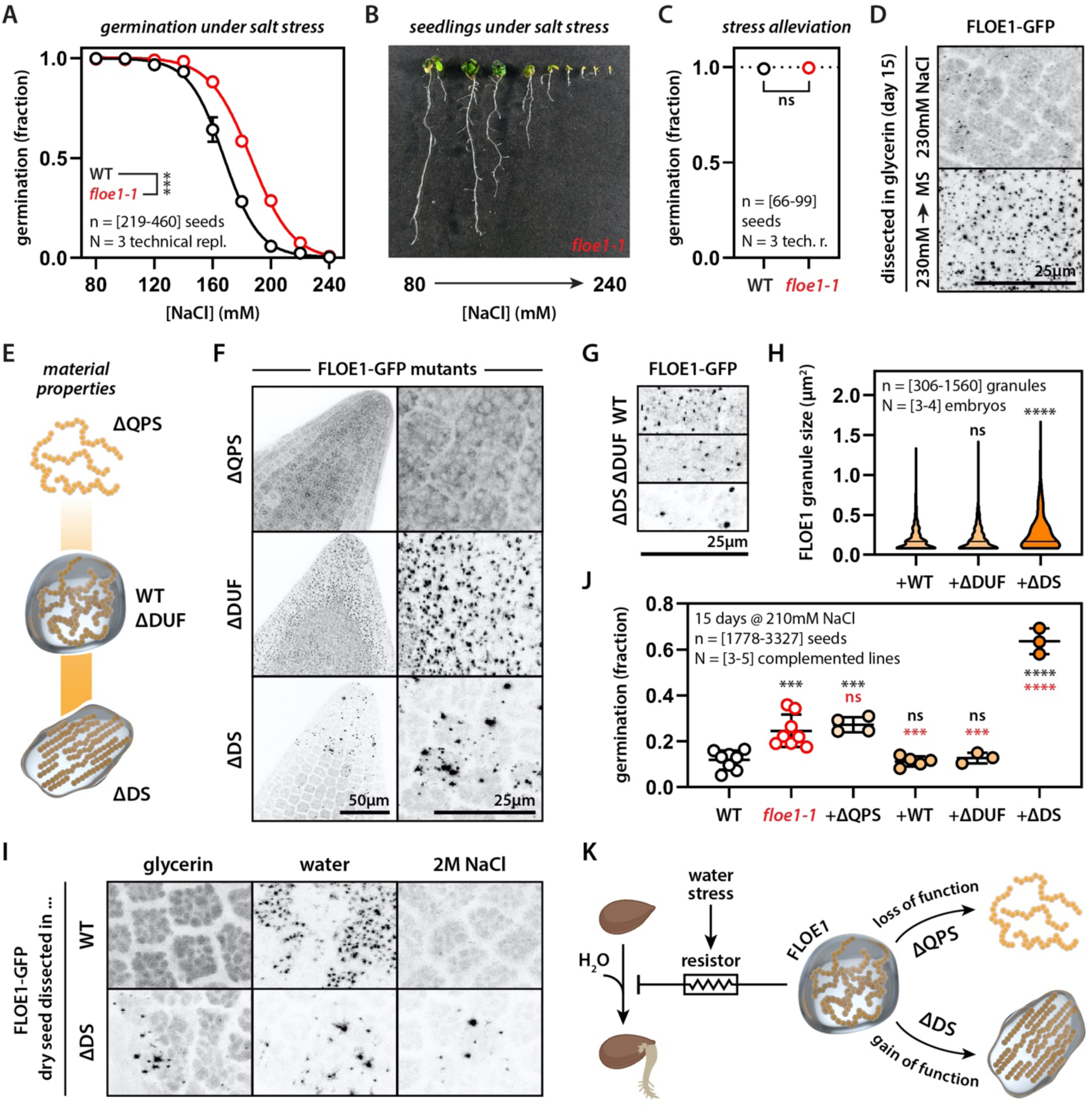
FLOE1 condensate material properties regulate its role in seed germination under salt stress. (A) *floe1-1* seeds show higher germination rates under salt stress. Two-way ANOVA. Four-parameter dose-response fit. Representative of two independent experiments. (B) Seedlings show developmental defects under salt stress. Three-week-old *floe1-1* seedlings are shown (wildtype shown in Fig. S8G). (C) Seeds retain full germination potential under standard conditions after a 15-day 230mM salt stress treatment. Representative of two independent experiments. (D) FLOE1 condensates are largely absent in ungerminated seeds after 15 days of incubation under salt stress. FLOE1 condensates appear within two hours after transfer to standard conditions (MS medium). (E) Scheme highlighting position of tested FLOE1 mutants on the material properties spectrum. (F) Representative images of floe1-1 mutants complemented with ΔQPS, ΔDUF and ΔDS forms of FLOE1 upon dissection in water. Embryo radicles are shown. (G) Close-up images of WT and mutant FLOE1 condensates. Embryo radicles are shown. (H) Quantification of FLOE1 condensate size. One-way ANOVA. (I) ΔDS FLOE1 condensates are not dependent on hydration. Embryo radicles are shown. (J) Germination rate of WT, *floe1-1* and complemented lines. One-way ANOVA. Representative of three independent experiments. (K) Scheme highlighting FLOE1’s role in regulating germination and the effect of mutants with altered material properties. * p-value < 0.05, ** p-value < 0.01, *** p-value < 0.001, **** p-value < 0.0001.

To assess whether FLOE1 phase separation is conserved, we explored the diversity of this protein in nature. First, we found that *A. thaliana FLOE1* encodes two isoforms from alternatively spliced transcripts (Fig. 4A). Besides the full-length isoform (FLOE1.1), *FLOE1* encodes a shorter splice isoform that lacks the majority of the DS domain (Fig. 4A). This shorter isoform (FLOE1.2) forms larger condensates (Fig. 4B) that can recruit the longer isoform (Fig. 4C). Second, searching the *A. thaliana* genome, we found two *FLOE1* paralogs, *FLOE2* (AT5G14540) and *FLOE3* (AT3G01560), which also formed larger condensates when expressed in tobacco leaves (Fig. 4D). Third, broadening our search, we found FLOE homologs in all green plant lineages, even in those preceding seed evolution (Fig. 4E-F, Fig. S10-11). Phylogenetic analysis revealed the emergence of two major clades (FLOE1-like and FLOE2-like), which show conserved variation in the length of the two disordered domains (Fig. 4G). By testing FLOE homologs across the plant kingdom, we have provided evidence for phenotypic variation in phase separation (Fig. 4H, Fig. S10-11) that is strikingly concordant with our engineered FLOE1 mutants (Fig. 2C).

**Figure 4:**
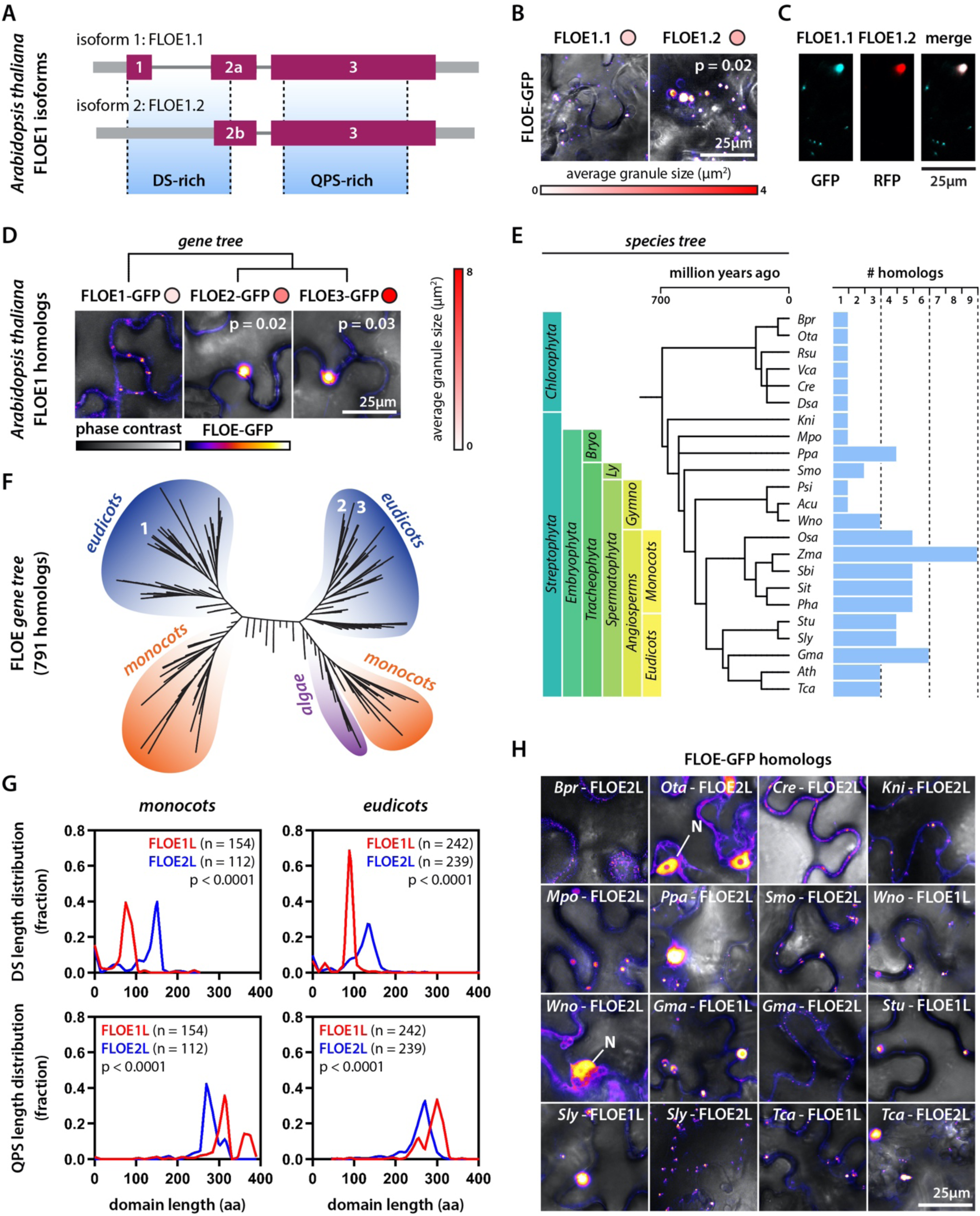
Natural sequence variation tunes FLOE phase separation. (A-B) *Arabidopsis thaliana* has two FLOE1 isoforms. FLOE1.2 is missing most of the DS-rich region (A) and has larger condensates than FLOE1.1 in tobacco leaves (B). [446-459] granules over 3 biological replicates. Mann-Whitney. (C) The large FLOE1.2 condensates recruit FLOE1.1. (D) FLOE1 has two *A. thaliana* paralogs that form larger condensates in tobacco leaves. [35-172] granules over 3 biological replicates. Mann-Whitney. (E) A species tree of the plant kingdom with example species and their number of FLOE homologs. (F) Gene tree of FLOE homologs. Numbers correspond to *A. thaliana* FLOE1, FLOE2 and FLOE3. (G) Distribution of DS and QPS length differences between the FLOE1-like and FLOE2-like clades among monocots and eudicots. Mann-Whitney. (H) Examples of FLOE homologs from across the plant kingdom. N denotes nuclear localization. For full species names for (E,F) see Fig. S10.

Phase separation is emerging as a universal mechanism to explain how cells compartmentalize biomolecules. Recent work in yeast suggests that phase separation of prion-like and related proteins is important for their function (*24, 25*), but this picture is less clear for multicellular organisms, especially since aggregation of these proteins is implicated in human disease (*26*). There is evidence suggesting the functionality of prion-like condensates in plants (*27–29*) and flies (*30*), but *in vivo* evidence for a functional role of the emergent properties of phase separation remains lacking. Here we demonstrate that conformational switches between liquid and solid-like states of FLOE1 can drive functional phenotypic variability via bet-hedging in a multicellular organism, similar to yeast (*14, 25*). Plant seed germination follows a bet-hedging strategy by spreading the risk of potentially deleterious conditions, such as drought, across different phenotypes in a population (*21–23*). Our data show that altering FLOE1’s material properties can tune these strategies in different environments. While the exact molecular mode of action of this newly discovered protein is still unclear, RNA-seq analysis suggests that its function is upstream of key germination pathways in a stress-dependent manner (Fig. S12 A-B, Table S2). We hypothesize that FLOE1 might act as a molecular glue helping to stabilize the desiccated glassy state (*31*), a model supported by an age-dependent loss of germination potential for *floe1-1* seeds (Fig. S12C). This also would suggest that the reversibility of FLOE1 condensation between the dry and the imbibed state is important for its function, which is consistent with the gain-of-function phenotype we observed with the irreversible DS mutant. While FLOE1 is the first reported protein to undergo hydration-dependent phase separation, it is likely that similar processes occur in a wide variety of organisms with quiescent desiccated life stages, including human pathogens, nematodes, fungal spores (*32–34*). Moreover, the large repertoire of FLOE sequence variation in the plant lineage suggests the possibility that natural populations may have used phase separation to fine-tune biological function to their ecological niches. Lastly, our work provides a set of guiding principles that can inspire the development of designer crops engineered to withstand the detrimental effects of climate change.

## Supporting information

Table S1. Seed Proteome

Table S2. RNA-seq analysis

Table S3. Sequences

## ACKNOWLEDGEMENTS

We thank Drs. M.B. Mudgett, D. Jarosz, H. Meyer, B. Schmidt, K. Lasker, C. Cuevas-Velasquez, and members of the Gitler and Rhee laboratories as well as the Carnegie-Stanford Intrinsically Disordered Protein Scientific Interest Group (IDPSIG) for helpful discussion and suggestions. We are grateful to Drs. Z. Wang’s and T. Nakagawa’s labs for sharing reagents, G. Materassi-Shultz for growth facilities management, and Dr. N. Boruah for bioinformatics assistance.

## FUNDING

This research was supported, in part, by the U.S. Department of Energy, Office of Science, Office of Biological and Environmental Research, Genomic Science Program grant nos. DE-SC0018277, DE-SC0008769, and DE-SC0020366 and by U.S. National Science Foundation grant MCB-1617020. **S.B.** acknowledges an EMBO Long Term Fellowship. Work in the **A.D.G.** lab is supported by NIH (grant R01 R35NS097263).

## AUTHOR CONTRIBUTIONS

**Y.D.:** Conceptualization, Methodology, Software, Validation, Formal Analysis, Investigation, Data Curation, Writing-Original Draft, Visualization. **S.B:** Conceptualization, Methodology, Validation, Formal Analysis, Investigation, Data Curation, Writing-Original Draft, Visualization. **B.J.:** Investigation, Validation. **F.B.:** Investigation**. E.F.:** Investigation, Methodology. **E.L.:** Investigation. **E.M.:** Investigation. **M.DD.:** Investigation. **P.B.:** Investigation. **A.S.H.:** Software, Writing-Review & Editing. **S.S.:** Supervision, Methodology, Writing-Review & Editing. **A.D.G.:** Conceptualization, Supervision, Funding acquisition, Writing-Review & Editing. **S.Y.R.:** Conceptualization, Supervision, Funding acquisition, Writing-Review & Editing.

## COMPETING INTERESTS

The authors declare no competing interests.

## DATA AND MATERIALS AVAILABILITY

All data is available in the manuscript or the supplementary materials. Seeds and constructs are available upon request.

## MATERIALS & METHODS

### Identification and analysis of the seed proteome

*Arabidopsis thaliana* genes were scored via the Expression Angler tool based on similarity to a “Developmental Map” expression pattern with “High Relative Expression” in “Dry Seed” and “Low Relative Expression” for all other tissues (https://bar.utoronto.ca/ExpressionAngler/) (*1*). The output was then normalized to Z-scores (Table S1) and genes were considered as seed-specific if they had a Z score of 3 or higher. The MobiDB-lite disorder scores of each gene in the “Z > 3” and “Z < 3” groups were retrieved from the MobiDB (version 3.1) *A. thaliana* dataset (https://mobidb.bio.unipd.it/dataset) (*2*), and their amino acid compositional profiles were obtained using the protr package (version 1.6-2) (*3*) in R. Genes in the “Z > 3” group were then checked for the presence of a predicted prion-like domain based on a list of Arabidopsis prion-like proteins (*4*). For FLOE1’s disorder prediction per residue, we used PONDR VSL2 (http://www.pondr.com) (*5*) and for identifying its prion-like domain we used PLAAC (http://plaac.wi.mit.edu) (*6*).

### Plant growth conditions

*Arabidopsis thaliana* plants from which seeds were harvested for the experimental assays (except for those used in the seed aging experiment) were grown in soil (PRO-MIX® HP Mycorrhizae) inside growth cabinets (Percival) held at 22 °C and 55 % humidity with a 16/8 hour photoperiod (130 μmol.m^−2^.s^−1^). Seeds were stratified for 3 days at 4 °C in the darkness to break dormancy. Plants from each line were randomly distributed and rotated every day until bolting to minimize environmental variations. When siliques began to mature, humidity was decreased to 45% as recommended by the Arabidopsis Biological Resource Center (ABRC) (ftp://ftp.arabidopsis.org/ABRC/abrc_plant_growth.pdf). Harvested seeds were air-dried for a week before being stored in Eppendorf tubes at 4 °C.

*Arabidopsis thaliana* plants that were used for line propagation and from which seeds were harvested for the seed aging experiments were grown in soil (PRO-MIX® HP Mycorrhizae) inside chambers held at 22 °C with a 16/8 hour photoperiod (130 μmol.m^−2^.s^−1^). Seeds were stratified for 3 days at 4 °C in the darkness to break dormancy.

*Nicotiana benthamiana* plants were grown in soil (PRO-MIX® PGX) inside chambers held at 22 °C with a 16/8 hour photoperiod (130 μmol.m^−2^.s^−1^).

In this study, “MS medium” refers to Murashige and Skoog medium with the following recipe: 0.5X Murashige and Skoog basal salt mixture (PhytoTechnologies Laboratories) at a pH of 5.7 supplemented with 0.8 % agar (Difco) and 1 % sucrose (Sigma-Aldrich)).

### Plant material

#### floe1-1 T-DNA mutant

The mutant line *floe1-1* (SALK_048257C) was obtained from ABRC and genotyped using primers priFLOE1cds-FWD/REV and the Salk genotyping primer LBb1.3 (Table S3). It was confirmed to be a knockout mutant by RT-qPCR (Fig. S8D) as described in the *RT-qPCR analyses* section.

#### Transgenic Lines

Transgenic plants were generated by *Agrobacterium tumefaciens*-mediated (GV3101 strain) transformation (*7*) of *floe1-1* with the constructs described in the *Plant plasmid construction* section, with the exception of the control transgenic line overexpressing YFP-FLAG used in Fig. S2A that was generated by introducing the transgene into Col-0. Transgenic seedlings (T_1_) were selected with Basta and T_2_ lines containing only one T-DNA construct were selected for further characterization by determining the Mendelian segregation ratio (3:1) of Basta-resistant seedlings in their progeny. Homozygote T_2_ lines were then identified by verifying that T_3_ seedlings were all Basta-resistant.

#### CRISPR lines

FLOE1 CRISPR lines were generated using the *Staphylococcus aureus* CRISPR-Cas9 system (*8*) and by following the protocol described in (https://www.botanik.kit.edu/molbio/940.php). A region within the QPS-rich region was identified as having a NNGGGT protospacer adjacent motif (PAM) downstream of a protospacer sequence (5’ TTACAGCCCCCAGACTGGC 3’) that did not have any significant similarities to other genomic regions. The corresponding guide RNA was inserted in the BbsI site of the pEn-Sa-Chimera vector through digestion-ligation following hybridization of the oligo duplex priCRISPR-FWD/REV (Table S3). The resulting sgRNA coding vector was then transferred to pDe-Sa-CAS9 through LR (Thermo Fisher Scientific) recombination. The final binary destination vector was then used to transform *Agrobacterium* (GV3101 strain), which was used to transform Col-0 plants using the floral dip method (*7*). Seeds obtained from the T_0_ parental lines were sown on MS medium supplemented with 30 mg/L Kanamycin (G-Biosciences) for selection of successfully transformed transgenics. Selected T_1_ seedlings were then transferred to soil to mature. Genomic DNA was extracted from mature rosette leaves of each of these T_1_ plants and the Cas9-recognition site within *FLOE1* was amplified through PCR with Phusion DNA polymerase (Thermo Fisher Scientific) using primers prigenoCRISPR-FWD/REV (Table S3). Sequencing (Sequetech Inc.) of the amplicons revealed that twelve plants demonstrated heterogenous sequences at the targeted region, which were subsequently selected for growing the T_2_ generation. For each selected T_1_ plant, eight T_2_ progeny were grown, and PCR amplification followed by sequencing of the *FLOE1* amplicon was again performed on genomic DNA extracted from mature rosette leaves. Four individuals from this T_2_ generation (*floe1-2*, *floe1-3*, *floe1-4*, *floe1-5*) presented different homozygous mutations in the *FLOE1* amplicon (Table S3), leading to frameshift mutations and pre-mature stop codons in the QPS region, and were selected for further assays.

### Plant plasmid construction

Constructs were generated using the Gateway system (Thermo Fisher Scientific) and vectors from the pGWB601-661 collection (*9*) as follows:

#### Transgenes for Arabidopsis experiments

FLOE1’s genomic region spanning its promoter, as predicted by AGRIS (*10*), to its last coding codon was amplified by Phusion (Thermo Fisher Scientific) PCR from Col-0 DNA (extracted with DNeasy Plant Mini Kit (Qiagen)) using the prigFLOE1-FWD/REV primers (Table S3). The amplicon was first cloned into pDONR221 (Thermo Fisher Scientific) using BP Clonase II (Thermo Fisher Scientific) and then subcloned into pGWB604, pGWB610 and pGWB633 (*9*) using LR Clonase II (Thermo Fisher Scientific) to generate pFLOE1p:FLOE1-GFP, pFLOE1p:FLOE1-FLAG and pFLOE1p:FLOE1-GUS respectively.

pFLOE1p:FLOE1ΔDS-GFP, pFLOE1p:FLOE1ΔQPS-GFP, and pFLOE1p:FLOE1ΔDUF-GFP were obtained by modifying pFLOE1p:FLOE1-GFP using the Q5 Site-Directed Mutagenesis Kit (New England Biolabs) with primers priDSdeletion-FWD/REV, priQPSdeletion-FWD/REV, and priDUFdeletion-FWD/REV (Table S3) respectively.

An entry vector containing the *YFP* gene was donated by Dr. Zhiyong Wang (Carnegie Institution for Science, USA) and another one, G18395, containing *FLOE1*’s coding sequence was obtained from ABRC. The two genes were then transferred from the entry vector into the binary vector pB7HFC3_0 (*11*) using Gateway cloning (Life Technologies), to create the vectors p35S:YFP-FLAG and p35S:FLOE1-FLAG.

Transgenes for tobacco (*Nicotiana benthamiana*) experiments:

A. *Arabidopsis* genes: The coding sequences of *FLOE1*’s isoforms, *FLOE1.1* and *FLOE1.2*, were amplified by Phusion (Thermo Fisher Scientific) PCR from the entry vector G18395 using priFLOE1.1-FWD/REV and priFLOE1.2-FWD/REV (Table S3) and then BP recombined into pDONR221 (Thermo Fisher Scientific). These were then transferred by LR recombination into pGWB605 (*9*) to generate p35S:FLOE1.1-GFP and p35S:FLOE1.2-GFP. Similarly, p35S:FLOE1.2-RFP was generated by subcloning FLOE1.2 into pGWB660 (*9*). The N-terminal version p35S:GFP-FLOE was generated by LR recombination of G18395 into pGWB606 (*9*). To generate p35S:FLOE2-GFP and p35S:FLOE3-GFP, the coding sequences of *FLOE2* and *FLOE3* were obtained from cDNA from 5-day old Col-0 seedlings by PCR amplification using Phusion DNA polymerase (Thermo Fisher Scientific) and the primers priFLOE2 -FWD/REV and priFLOE3-FWD/REV (Table S3). Total cDNA was obtained by reverse transcription using M-MLV Reverse Transcriptase (Thermo Fisher Scientific) from total RNA extracted with the RNeasy Plant Mini Kit (Qiagen). The *FLOE2* and *FLOE3* amplicons were then BP recombined into pDONR221 before being transferred into pGWB605 by LR recombination to generate p35S:FLOE2-GFP and p35S:FLOE3-GFP.
B. Mutated FLOE1 versions: FLOE1wt, FLOE1Δnucl, FLOE1ΔCC, FLOE1ΔQPS, and FLOE1-QPS-15xY-S were amplified from the corresponding human expression vectors described in *Human plasmid construction* using prihFLOE1-FWD/REV (Table S3), and BP recombined into pDONR221 (Thermo Fisher Scientific) before being transferred by LR recombination into pGWB605 to generate p35S:FLOE1wt-GFP, p35S:FLOE1Δnucl-GFP, p35S:FLOE1ΔCC-GFP, p35S:FLOE1ΔQPS-GFP, and p35S:FLOE1-QPS-15xY-S-GFP. p35S:FLOE1ΔDS-GFP and p35S:FLOE1ΔDUF-GFP were obtained by the same process but with different primer pairs: prihFLOE1ΔDS-FWD/prihFLOE1-REV and prihFLOE1-FWD/prihFLOE1ΔDUF-REV (Table S3), respectively.
C. Non-*Arabidopsis* FLOE1 homologs: Protein sequences for all FLOE1 homologs shown in Fig. 4H and Fig. S10B were obtained from UniProt (*12*) and Phytozome v13 (*13*). Their corresponding DNA sequences were generated with codon-optimization for *Nicotiana benthamiana* expression using IDT’s codon optimization tool (https://www.idtdna.com/CodonOpt) (Table S3). The sequences were synthesized by GenScript Biotech Corporation (Piscataway, NJ) with flanking attB sites for subsequent BP cloning into pDONR221 (Thermo Fisher Scientific). They were then subcloned into pGWB605 by LR recombination to generate p35S:HOMOLOG-GFP constructs (where HOMOLOG refers to the relevant FLOE1 homolog).

### Analysis of FLOE1 homologs

#### Phylogenetic tree construction

All *Viridiplantae* protein sequences containing the highly-conserved DUF1421 domain were retrieved from UniProt (*12*). After removal of duplicates due to re-annotations, the remaining 791 sequences (Table S3) were submitted to the phylogenetic analysis tool NGPhylogeny.fr (*14*) with default settings. The FastME Output Tree was then uploaded to iTOL (version 5) (*15*) for tree visualization.

#### Lengths of QPS and DS domains

All monocot and eudicot sequences from the FLOE1-like and FLOE2-like clades were aligned using the msa package (version 1.20.0) in R (*16*). The DS and QPS regions of the homologs were defined as aligning to the DS and QPS regions of FLOE1. The lengths of these regions were used for subsequent analysis.

#### Multiple Sequence Alignments

Alignments and protein characteristics displayed in Fig. S11 were conducted using the msaPrettyPrint() function of the msa package (*16*) and using TEXshade (version 1.25) (*17*) in R and MacTex.

### Tobacco infiltration

*Agrobacterium* cultures (GV3101 strain) carrying the relevant constructs were grown overnight at 28 °C in LB broth (Fisher BioReagents) containing 25 mg/L rifampicin (Fisher BioReagents), 50 mg/mL gentamicin (GoldBio) and 50 mg/L spectinomycin (GoldBio). Cultures were washed four times with infiltration buffer (10 mM MgCl_2_ (omniPur, EMD), 10 mM MES (pH 5.6) (J. T. Baker) and 100 uM acetosyringone (Sigma-Aldrich)) and diluted to reach an OD_600_ of 0.8. Fully expanded 3^rd^, 4^th^, or 5^th^ leaves from 6-week-old tobacco plants were infiltrated with these diluted *Agrobacterium* cultures using Monoject 1mL Tuberculin Syringes (Covidien). For the FLOE1.1-GFP and FLOE1.2-RFP colocalization experiment, an equal amount of each culture was pre-mixed before infiltration. For each construct or combination of constructs, at least three individual tobacco plants were infiltrated.

### Germination experiments

Seeds were first sterilized by vortexing in 70 % ethanol for 5 minutes after which the solution was removed and replaced with 100 % ethanol. Seeds were then placed on pre-sterilized filter papers (Grade 410, VWR) and left to dry in a laminar flow hood. Sterilized seeds were then sown on square petri dishes (120 x 120 wide x 15 mm high (VWR)) containing 40 mL of MS medium (See *Plant growth conditions* section) supplemented with NaCl (Sigma-Aldrich) and mannitol (Sigma-Aldrich) at the concentrations indicated below. Plates were then sealed with micropore surgical tape (3M) and covered in aluminum foil before being placed at 4°C. After exactly 120 h (5 days) of stratification to break seed dormancy, plates were transferred to a 24 h light (130 μmol.m^−2^.s^−1^), 22 °C growth cabinet (Percival). Germination (identified by radicle protrusion) was counted under a dissecting microscope.

Germination experiments were performed on seeds from three independent batches of plants (A, B, and C) grown as described in the *Plant growth conditions* section. Germination data displayed in this publication are representative of 2-3 independent experiments as indicated in the relevant legends.

#### Batch A (Fig. 3, Fig. S8E-H, Fig. S9)

40 Col-0 and 40 *floe1-1* plants were grown alongside 10 plants of each of the following lines: four independent CRISPR lines (*floe1-2, floe1-3, floe1-4, floe1-5*), five independent pFLOE1p:FLOE1-GFP lines, two independent pFLOE1p:FLOE1-FLAG lines, one pFLOE1p:FLOE1-GUS line, three independent FLOE1p:FLOE1ΔDS-GFP lines, four independent FLOE1p:FLOE1ΔQPS-GFP lines, and three independent FLOE1p:FLOE1ΔDUF-GFP lines. For each line, seeds from five plants were randomly pooled together which resulted in 2 biological replicates of each CRISPR and complemented line, and 8 biological replicates of Col-0 and *floe1-1.* For each biological replicate and each germination condition (0, 80 mM, 100 mM, 120 mM, 140 mM, 160 mM, 180 mM, 195 mM, 200 mM, 210 mM, 220 mM, 230 mM and 240 mM NaCl), three technical replicates were conducted. At the end of the 230 mM NaCl germination experiment (day 15), seeds that did not germinate were rinsed in sterile double distilled water and sown on normal MS medium. Two days later, germination was scored to test whether they maintained their germination potential. The 80mM, 100mM, 120mM, 140mM, 160mM, 180mM, 200mM, 220mM, and 240mM NaCl experiments were conducted independently from the 195mM, 210mM and 230mM NaCl experiments.

#### Batch B (Fig. S8A-D)

14 Col-0 and 27 *floe1-1* plants were grown alongside 6 plants of each of the following lines: three independent pFLOE1p:FLOE1-GFP lines, two independent pFLOE1p:FLOE1-FLAG lines, one pFLOE1p:FLOE1-GUS line, and two independent 35S:FLOE1-FLAG lines. The 35S:FLOE1-FLAG lines failed to express *FLOE1* as revealed by RT-qPCR (Fig. S8D) and were therefore chosen as transgenic controls. Seeds from each individual plant were sown on MS medium (See *Plant growth conditions* section) supplemented with either mannitol (400 mM) or NaCl (190 mM, 205 mM and 220 mM). For each biological replicate and each germination condition, three technical replicates were conducted.

#### Batch C (Fig. S12C)

5 *floe1-1* plants and 5 Col-0 plants were alternated within the same flat. Seeds from each individual plant were harvested and aged in Eppendorf tubes placed inside an opaque box stored at room temperature for 42 months (3.5 years). They were then sown on MS medium (See *Plant growth conditions* section). For each biological replicate, three technical replicates were conducted.

### Embryo dissection and *in vivo* FLOE1 visualization experiments

#### Salt, mannitol, sorbitol, cycloheximide and water assays

Seeds of the relevant GFP-tagged lines were submerged in either glycerin or in solutions of NaCl (Sigma-Aldrich), mannitol (Sigma-Aldrich), sorbitol (Sigma-Aldrich), cycloheximide (GoldBio) or double distilled water at concentrations indicated in the publication for 15-30 min (NaCl: 0, 0.2 M, 0.4 M, 0.6 M, 0.8 M, 1 M, 1.2 M, 1.4 M, 1.6 M, 1.8 M, 2 M; mannitol: 0, 0.95 M; sorbitol: 0, 0.725 M, 1.45 M; cycloheximide: 1 g/L). They were then dissected to remove the seed coat and imaged by confocal microscopy (see *Plant live imaging microscopy and image analysis* for details). As controls, 35S:GFP (*11*) seeds were dissected in water to verify that GFP alone could not induce condensate formation, and Col-0 seeds were dissected in either water or 2 M NaCl to assess the level of autofluorescence of the protein storage vacuoles in the absence of GFP in these conditions.

#### Condensate reversibility assays

Three different types of FLOE1 condensate reversibility assays were performed: 1) Embryos from dry seeds were first dissected in glycerin as described above, and after imaging, glycerin was washed off from the embryos with water and the same embryos were imaged immediately in water (less than 5 minutes); 2) Seeds were submerged in water for 1 h before being transferred to 2 M NaCl for 10 min and imaged and *vice versa* (1 h in 2 M NaCl followed by 10 min in water); and 3) Seeds were submerged in water overnight and then left to dry for an additional day. Seeds were then either dissected in glycerin to obtain the condensate state of the dry seeds or in water to assess the ability to re-form condensates.

#### End of germination experiment analysis

At the end of the 230 mM NaCl germination experiment described in the *Germination experiments* section (5 days of stratification followed by 15 days in light on MS medium supplemented with 230 mM NaCl), seeds that did not germinate were either: 1) dissected directly in glycerin to leave the hydration state of the seed unaltered; or 2) transferred first to standard MS medium and dissected in glycerin two hours later. Dissected embryos were then imaged by confocal microscopy to obtain a snapshot of their final condensate state (see *Plant live imaging microscopy and image analysis*).

#### Developmental stages of the embryos

FLOE1p:FLOE1-GFP and 35S:YFP-FLAG flower buds were self-crossed 11, 8, 6 and 4 days before dissection to obtain developing siliques carrying embryos at mature, torpedo, heart and globular stages respectively. Seeds from the various developmental stages were dissected either in glycerin or water and imaged by confocal microscopy (see *Plant live imaging microscopy and image analysis*).

### GUS staining

FLOE1p:FLOE1-GUS seeds carrying embryos at different developmental stages were incubated at 37 °C overnight in GUS staining solution (*18*). In the case of dry seeds, seed coats were first removed (as they were impermeable to the staining solution) and embryos were incubated at 37 °C for one hour in GUS staining solution. Following the incubation, samples were destained in 70 % ethanol at room temperature for 24 hours and embryos were dissected out (in the case of developing siliques) before imaging. Pictures were taken with a compound microscope (Nikon) and a dissecting microscope (Leica MZ6 microscope).

### Plant live imaging microscopy and image analysis

#### Image acquisition

Embryos and tobacco leaves were imaged at room temperature on a LEICA TCS SP8 laser scanning confocal microscope in resonant scanning mode using the LASX software. All samples were imaged with a HC PL APO CS2 63X/1.20 water objective with the exception of embryos submerged in glycerin that were imaged with a 63X/1.30 glycerin objective and of embryos of early developmental stages that were imaged with a HC PL APO CS2 20x/0.75 dry objective. GFP, RFP, or YFP fluorescence was detected by exciting with a white light laser at 488 nm, 561 nm and 514 nm, respectively, and by collecting emission from 500-500 nm, 591-637 nm and 524-574 nm, respectively, on a HyD SMD hybrid detector (Leica) with a lifetime gate filter of 1-6 ns to reduce background autofluorescence due to chlorophyll (tobacco) or protein storage vacuoles (embryos). Z-stacks were collected with a bidirectional 96-line averaging while single-frame images (tobacco images displayed in the publication) were collected with a bidirectional 1024-line averaging. With the exception of the single-frame embryo images shown in Fig. 1F and Fig. 3G, all embryo images shown in this publication are maximum intensity projection images. For the colocalization experiments, samples were imaged sequentially between each line to ensure that the colocalization signals were not due to bleed-throughs. Images displayed in the publication were representative of at least three biological replicates for each construct (tobacco) or line (Arabidopsis). All samples that were compared in the publication were imaged with the same magnification and laser intensity.

#### Intracellular heterogeneity analysis

For each radicle and experimental condition, maximum projection images of their corresponding Z-stacks were obtained using the LASX software. ROIs were then manually drawn around each individual cell to obtain their standard deviation (RMS) and mean intensity levels. Heterogeneity scores were obtained by dividing the standard deviation by the mean (coefficient of variation). Between 363 and 461 cells were measured per embryo with a total of 3 embryos per condition. Cells were characterized as exhibiting FLOE1 condensates if their heterogeneity score was higher than the top 5 percentiles of the 2 M NaCl condition (heterogeneity cut-off = 0.3 arbitrary units (a.u.)).

#### FLOE1 condensate size determination

Individual slices of an embryo radicle Z-stack were analyzed using FIJI (*19*). Individual condensates were identified using a threshold, and subsequently measured for their area. A total of 3-4 embryos per condition were analyzed.

### Seed phenotyping

#### Seed weight

Seeds from twelve and fourteen biological replicates of *floe1-1* and Col-0, respectively, were used for the seed weight analysis. Seeds were weighed on a Sartorius M2P scale in batches of nine to twenty seeds and the process was replicated three times per biological replicate. The average weight per seed was calculated and used for subsequent statistical analysis.

#### Seed size and aspect ratio

Seeds from fourteen and sixteen biological replicates of *floe1-1* and Col-0, respectively, were used for the seed size and aspect ratio analyses. Seed images were scanned using a Canon CanoScan LiDE 700 F (Canon Inc). All images were scanned at 600 dpi and, for ease of collection, the seeds were placed in transparent bags before scanning. The number of seeds per image varied, but ten seeds per sample were randomly selected and analyzed for area quantification and aspect ratio using ImageJ (version 2.0.0) (*20*). This process was replicated ten times per biological replicate to obtain a total of one hundred seeds per biological replicate.

### RNA extraction from seeds and siliques

DNA-free total RNA was extracted from seeds and siliques as described in (*21*). The extraction buffer utilized 0.5% β-mercaptoethanol. RNA quantity and purity from all samples were assessed using a NanoDrop Spectrophotometer (Thermo Fisher Scientific).

### RT-qPCR analyses

cDNA was synthesized from 1 μg of extracted RNA using M-MLV Reverse Transcriptase (Thermo Fisher Scientific), per manufacturer’s protocol. qPCR was performed using the SensiFAST SYBR No-ROX Kit (Bioline). Primers used to quantify *FLOE1* expression were priqPCRFLOE1set1-FWD/REV (Table S3), with the exception of the qPCRs conducted on the CRISPR lines (Table S3) as well as on siliques and seeds from different developmental stages (Fig. S1B) where priqPCRFLOE1set2-FWD/REV were used (Table S3). The different developmental stages of siliques were defined based on color: dark green, light green and yellow, which roughly correspond to 4-7, 8-10 and 11-13 days post-anthesis, respectively (*22*). The reference gene that was used to normalize gene expression levels, At5G25760 (*PEX4*), was chosen for consistent expression in seeds (*23*). We used a primer pair, priAT5G25760-FWD/REV (Table S3), which was reported in (*24*). Reactions were run on 96-well plates in the LightCycler® 480 Instrument II system and were repeated three times.

### RNA-seq experimental conditions and analysis

#### Experimental design

RNA-seq analysis was carried out on six conditions: 1) dry *floe1-1* seeds; 2) dry Col-0 seeds; 3) imbibed *floe1-1* seeds; 4) imbibed Col-0 seeds; 5) *floe1-1* seeds imbibed with 220 mM NaCl; and 6) Col-0 seeds imbibed with 220 mM NaCl. Three biological replicates corresponding to pooled seeds from 20 plants were performed per condition, with 50 mg of mature seeds used per biological replicate. For conditions (1) and (2), RNA was extracted directly from dry seeds using the protocol described in the *RNA extraction from seeds* section. For conditions (3) and (4), dry seeds were sown onto separate agar plates of MS medium conditions and cold-stratified for 5 days at 4°C in the dark. All plates were subsequently transferred to and held in a growth cabinet (Percival) for 4 hours under light (130 μmol.m^−2^.s^−1^) and 22°C. After the 4-hour incubation, imbibed seeds were scraped from each plate and transferred to a clean mortar and pestle and ground in liquid nitrogen. RNA was then extracted as described in the *RNA extraction from seeds* section. Conditions (5) and (6) were conducted in parallel and using the same experimental settings with the MS medium supplemented with 220 mM NaCl.

Samples from all biological replicates were first sent to the Stanford University Protein and Nucleic Acid Facility (Stanford, CA) for quantification and quality analysis using a 2100 Bioanalyzer (Agilent). Samples were then sent to Novogene Corporation Inc. (Sacramento, CA) for RNA-seq library preparation (250-300 bp insert cDNA library) and sequencing (2×150 bp paired-end reads on an Illumina Platform).

#### Analysis

Reads were mapped with HISAT2 to the *Arabidopsis thaliana* TAIR10 reference genome using the Galaxy (Version 2.1.0+galaxy5) web platform (https://usegalaxy.eu) (*25*). The resulting BAM files were then analyzed on R using the DESeq2 (*26*) and TxDB.Athaliana.BioMart.plantsmart28 (Bioconductor) packages. Genes with padj<0.05 were considered differentially expressed. Gene Ontology and KEGG enrichment of the differentially expressed genes was obtained using g:Profiler (https://biit.cs.ut.ee/gprofiler/gost) (*27*).

### All-atom modeling and structural prediction

To obtain structural insight into the previously uncharacterized nucleation domain of FLOE1, we combined a set of integrative homology modelling approaches with all-atom simulations (Fig. S5). We first leveraged a number of homology-modelling based tools to perform *de novo* structural prediction, including SWISS-MODEL and Phyre2 (*28, 29*). SWISS-MODEL strongly predicted the majority of the nucleation domain to consist of a trimeric coiled-coil, while in agreement Phyre2 predicted a single long helix with the appropriate exposed interfacial residues necessary for trimeric coiled-coil formation. The secondary structure prediction server PSI-PRED identified a number of overlapping residues predicted to be helical (*30*). Taken together, all structural modelling tools predict this region to be highly helical and to form higher-order oligomeric assemblies, providing a molecular explanation for its observed cellular function.

To provide further support for the predicted helical structure of the nucleation domain, we turned to all-atom simulations. Simulations were performed with the ABSINTH implicit solvent model and CAMPARI Monte Carlo simulation engine (*31*). We first generated an initial helical starting structure by applying a bias force to drive the protein into a single alpha helix. From this helical starting state, production simulations were run in which no bias was applied, and the system was able to evolve. Five independent simulations were performed with the reported helicity being the average of all five runs. Simulations were run using the ABSINTH implicit solvent model and abs_3.2_opls.prm parameters. Each simulation was run for 6.8 x 10^7^ Monte Carlo steps with the first 6.0 x 10^6^ steps discarded as equilibration. These simulations revealed a remarkably good agreement with the predictions from homology modelling tools (Fig. S5E) including a break in helicity identified in the homology modeling tools between residues 130 and 140. Taken together, distinct structural and biophysical approaches converge on a model in which the nucleation domain drives higher-order assembly as a multivalent trimeric coiled-coil domain.

### Recombinant FLOE1 expression, purification, and phase separation

FLOE1’s coding sequence was transferred from G18395 (see *Plant vector construction* section) into pDEST-HisMBP (Addgene plasmid # 11085) (*32*) using LR Clonase II (Thermo Fisher Scientific).

A sequence encoding the TEV recognition sequence, ENLYFQ, was then added between the MBP and FLOE1 coding sequences using the Q5 Site-Directed Mutagenesis Kit (New England Biolabs) with primers priTEV-FWD/REV (Table S3). BL-21(DE3) competent cells (Agilent) were transformed with the plasmid following supplier protocol and plated on LB agar plated with ampicillin selection overnight at 37 °C. Transformed cells were expressed in 6 L of LB at pH 7.4 with ampicillin selection. Expression was induced at OD_600_ of 0.6 using 0.5 M IPTG, and cells were left under shaking at 220 RPM and 16 °C for 19 h prior to collection.

Collected cells were spun down at 4 °C for 8 min and the supernatant was discarded. The pellet was resuspended with 20 mL lysis buffer (50 mM NaH2PO4, 0.5 M NaCl, pH 8) per 1 L of expression, and the resuspended cells were homogenized for 8 min (Emulsiflex homogenizer). The resulting cell lysate was spun down for 50 min at 20,000 g, and the supernatant was collected. Ni-NTA nickel beads (Qiagen) were equilibrated with lysis buffer, then loaded with the lysate and washed with 50 mL of wash buffer (50 mM NaH2PO4, 0.5 M NaCl, 20 mM Imidazole, pH 8) followed by 8 mL of Elution Buffer (50 mM NaH2PO4, 0.5 M NaCl, 250 mM Imidazole, pH 8) all done at 4°C. The eluent was collected, spun down to remove aggregates, and further purified with size exclusion chromatography (SEC) using a Superdex 200 16/60 column (GE). 5 mL of spun-down eluent was injected onto the column and ran at 0.5 mL per min at room temperature. Fractions were collected and the presence of the protein was verified using SDS-PAGE. The sample corresponding to Peak #1 was purified separately from Peak #2 (Fig. S3A). Differential interference contrast (DIC) microscopy images were taken from each peak. Peak #1 showed many small aggregates (not shown) whereas Peak #2 was largely devoid of aggregates (Fig. S3B).

Following SEC, FLOE1-MBP from Peaks #1 and #2 was cleaved using Tobacco Etch Virus (TEV) protease using a 1:50 w/w TEV to FLOE1-MBP1 ratio overnight at 16°C with shaking at 200 RPM. The cleavage reaction was confirmed with SDS-PAGE (Fig. S3C). DIC images of the mixture following cleavage confirmed the appearance of round droplets in Peak #2 (Fig. S3B), while Peak #1 showed the same aggregates that were present prior to cleavage (data not shown).

Four successive rounds of SEC showed almost identical results, indicating that the protein remained intact for at least 24 hours. SDS-PAGE also showed almost identical bands from corresponding collected fractions. Of note is the distinction in properties between Peak #1 and Peak #2. Peak #1 elutes at the void volume of the column, meaning it is larger than what can be resolved with Superdex 200 (> 200 kDa) (Fig. S3A). We see that Peak #1 produces a clean band off SEC but results in aggregation in both the pre- and post-cleaved FLOE1. Peak #2 elutes afterwards, implying it is closer to a monomeric state. It shows very little aggregation before cleavage, but displays the spherical droplets indicative of phase separation after removing the MBP with TEV (Fig. S3B-C). This behavior has been observed for several other LLPS-forming proteins, such as FUS (*33*).

### Human plasmid construction

FLOE1 and derived mutant constructs for expression in human cells were optimized for human expression (Table S3) and generated through custom synthesis and subcloning into the pcDNA3.1+N-eGFP backbone by Genscript (Piscataway, USA).

### Human cell culture and microscopy

U2OS cells (ATCC, HTB-96) were grown at 37 °C in a humidified atmosphere with 5 % CO_2_ for 24 h in Dulbecco’s Modified Eagle’s Medium (DMEM), high glucose, GlutaMAX + 10 % Fetal Bovine Serum (FBS) and pen/strep (Thermo Fisher Scientific). Cells were transiently transfected using Lipofectamine 3000 (Thermo Fisher Scientific) according to manufacturer’s instructions. Cells grown on cover slips were fixed for 24 h after transfection in 4 % formaldehyde in PBS. Slides were mounted using ProLong Gold antifade reagent (Life Technologies). Confocal images were obtained using a Zeiss LSM 710 confocal microscope. Images were processed using FIJI (*19*).

### FRAP measurements in human cells

U2OS cells were cultured in glass bottom dishes (Ibidi) and transfected with GFP-FLOE1 constructs as described above. After 24 h, GFP-FLOE1 condensates were bleached and fluorescence recovery after bleaching was monitored using Zen software on a Zeiss LSM 710 confocal microscope with incubation chamber at 37 °C and 5 % CO_2_. Data were analyzed as described previously (*34*). In brief, raw data were background subtracted and normalized using Excel, and plotted using GraphPad Prism 8.4.1 software.

### Standard transmission electron microscopy (TEM) in human cells

U2OS cells were transfected at 80% confluency in Petri dishes, transfected for 24 h (see above) and fixed with 2.5 % glutaraldehyde in 0.1 M Na-cacodylate buffer (pH 7.2). After washing in the same buffer three times, the cells were scraped and centrifuged at 200 x g. The pellet was resuspended in 1.5 % low melting point agarose (type VIIa, Sigma-Aldrich) in 0.1M Na-cacodylate buffer and pelleted for 1 min at 1000x g. After solidification on ice, the agarose-embedded cell pellet was cut in small blocks, incubated in 1 % osmium tetroxide for 2 h at room temperature, washed 3x in milli-Q water and dehydrated in a graded ethanol series until 100 % ethanol in steps of 5 minutes. Finally, following 2 washes with propylene oxide, the cells were infiltrated with epoxy resin (Agar 100, EMS, Hatfield, PA, USA) embedded in BEEM-capsules and cured for 48 h at 60°. Ultrathin sections (70 nm) were cut from the polymerized samples with a Leica ultracut UCT ultramicrotome (Leica, Vienna, AU), and post-stained with 4% uranyl acetate (SPI Supplies, Westchester, PA, USA) for 8 mins and Reynold’s lead citrate for 3 mins. Finally, the cells were observed and imaged at an acceleration voltage of 80 kV with a JEOL JEM1400 (Tokyo, JP) transmission electron microscope equipped with an 11 Mpx EMSIS Quemesa camera (EMSIS GmbH, Muenster, DE).

### Correlative Light and Electron Microscopy (CLEM) in human cells

For CLEM, plated cells were lightly fixed with 2% PFA and washed 3x with PBS. The cells were scraped and pelleted at 200x g after which they were resuspended in 20 % BSA and pelleted again at 200x g. The loosely packed cells in BSA were high-pressure frozen in a Leica Empact 2 high-pressure freezer (Leica, Vienna, Au), and submitted to a quick freeze-substitution protocol that preserves fluorescence (*35, 36*). Briefly, frozen samples were freeze-substituted in acetone containing 0.2 % uranyl acetate and 5 % H_2_O in a Styrofoam box on a rotating platform while temperature was allowed to rise to −50°C at which moment they were transferred to the Leica AFS2 automatic freeze-substitution apparatus equipped with a Leica Freeze Substitution Processor (FSP) processing robot (Leica, Vienna, AU). After total time elapsed between −80°C and −50°C amounted 1.5 h, samples were washed in acetone and infiltrated in Lowicryl HM20 resin (EMS, Hatfield, PA, USA), and finally polymerized at −50 °C by UV-light. Sections of 100 to 200 nm were cut with a Leica ultracut UCT ultramicrotome and deposited on an indium-tin oxide coverslip, and fluorescent images were taken with a Zeiss Axioplan light microscope (Zeiss, Oberkochen, DE). After this, the ITO-coverslip with sections was mounted to a support stub, placed inside the specimen chamber of a Zeiss Sigma scanning electron microscope (Zeiss, Oberkochen, DE), and imaged at 1.25 - 1.5 kV acceleration voltage with a Gatan backscattered electron detector. For correlation, fluorescent and electron images were overlaid using GIMP (GNU Image Manipulation Program).

### Statistical analysis

All data was analyzed using Graphpad Prism 8.4.1 and Excel. Statistical tests details are shown in figure legends.

## SUPPLEMENTAL FIGURES

**Figure S1:**
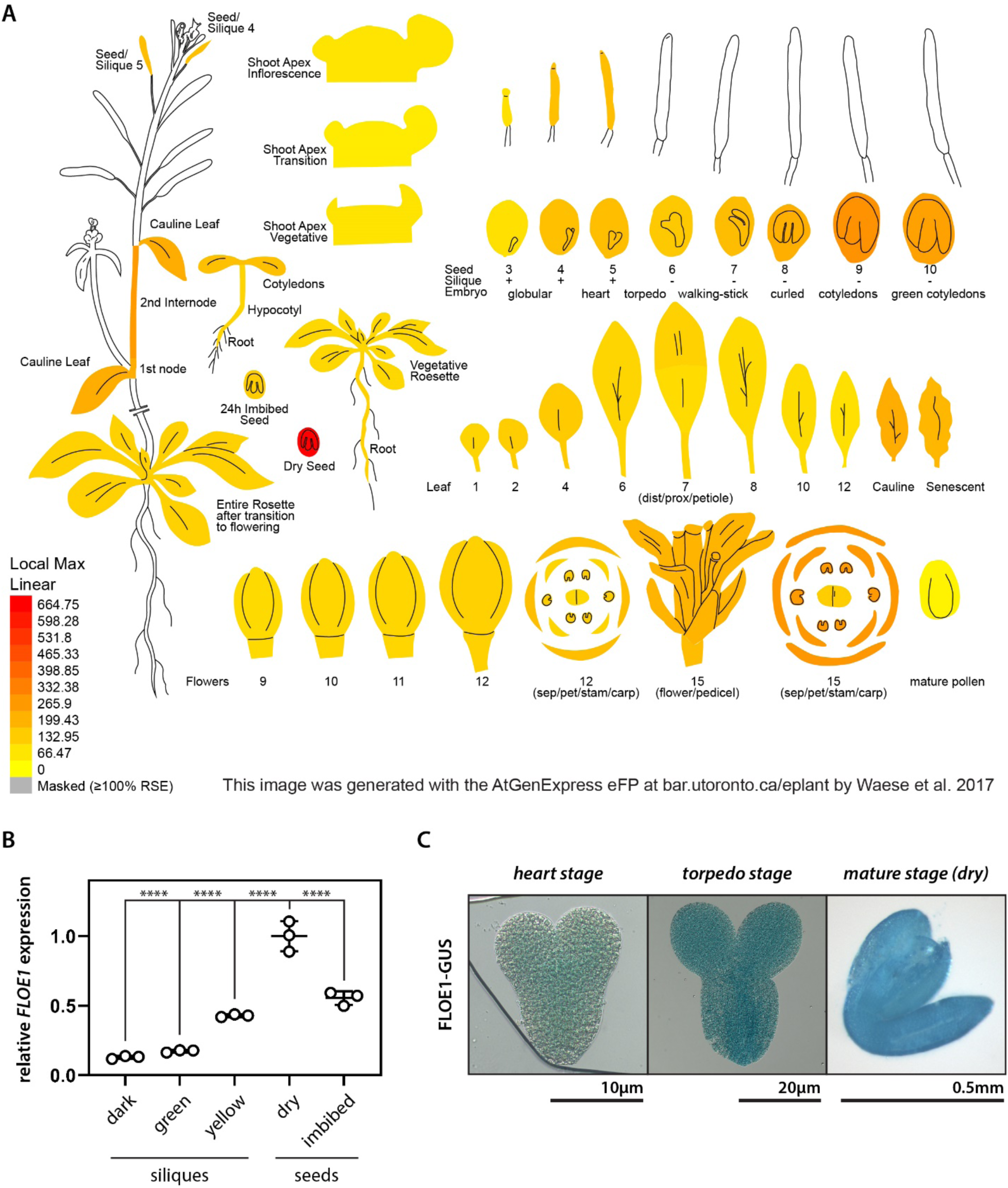
F*L*OE1 and FLOE1 expression in *Arabidopsis*. (A) Tissue-specific expression of *FLOE1* derived from ePlant (*37*). (B) RT-qPCR analysis of different developmental stages shows peak expression in mature dry seeds, and a decrease in expression upon imbibition. “Dark”, “green” and “yellow” refer to the maturation stages of the siliques (from younger to older), which roughly correspond to 4-7, 8-10 and 11-13 days post-anthesis (*22*), and “imbibed” corresponds to seeds that were imbibed in sterile double-distilled water for 24 h. Col-0 (WT) plants were used. One-way ANOVA. **** p-value < 0.0001. Mean ± SD shown. (C) Expression of FLOE1 in developing embryos detected by GUS staining in FLOE1p:FLOE1-GUS transgenic lines.

**Figure S2:**
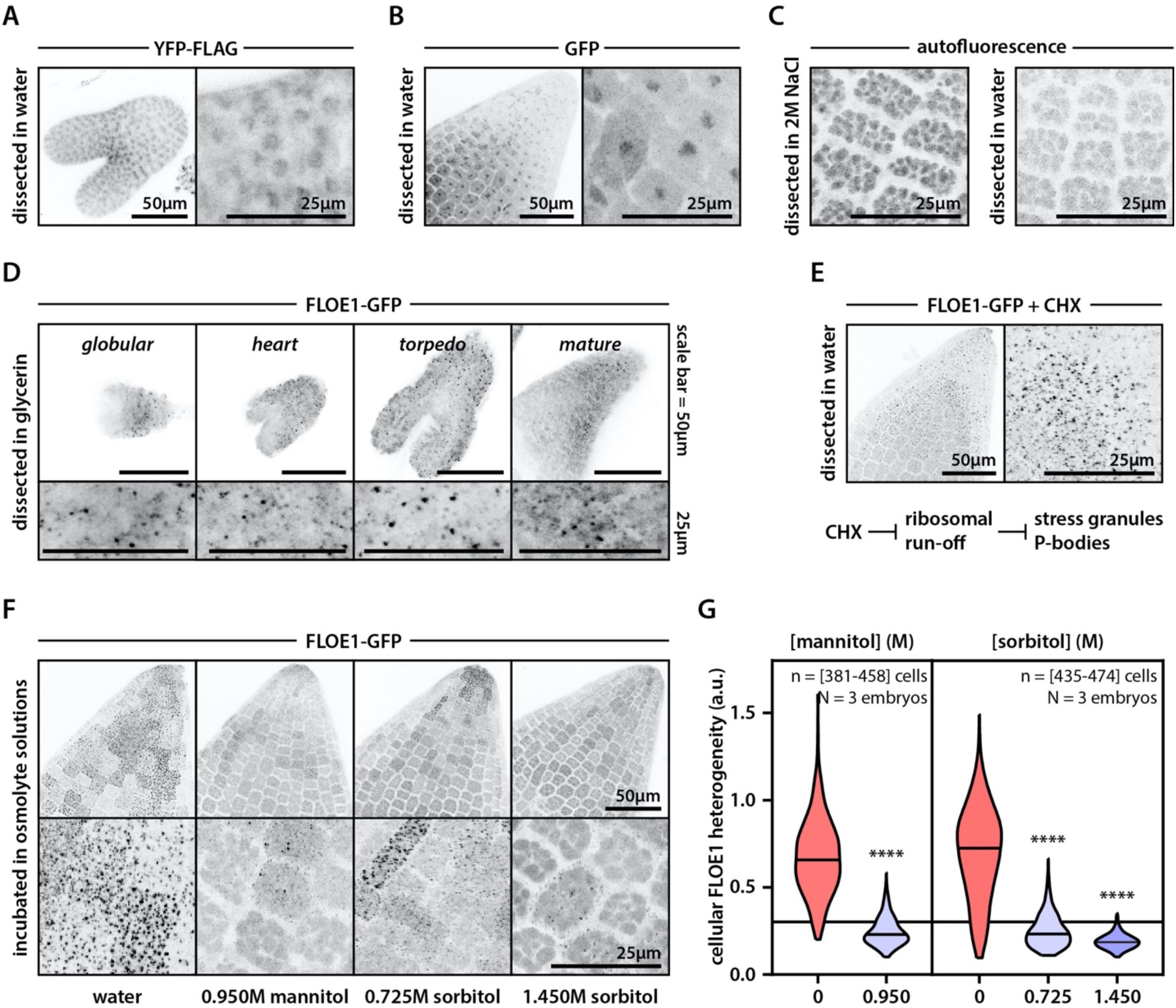
FLOE1 forms condensates dependent on the water potential of the cellular environment. (A) YFP-FLAG localizes diffusely with modest nuclear enrichment in *Arabidopsis* torpedo stage embryos without any condensates forming. (B) GFP localizes diffusely with modest nuclear enrichment in imbibed dry seed-derived embryo radicles without any condensates forming. (C) Autofluorescence of protein storage vacuoles in non-transgenic control plants is much weaker in the hydrated state. (D) Dissection in glycerin does not alter the presence of FLOE1-GFP condensates throughout embryonic development before desiccation. For the mature stage, a zoom in of the radicle is shown. (E) Cycloheximide (CHX) treatment does not prevent FLOE1-GFP condensate formation in imbibed embryo radicles. (F-G) Incubation of FLOE1-GFP embryos in osmolyte solutions prevents FLOE1 condensate formation. Mannitol: Mann-Whitney. Sorbitol: One-way ANOVA Kruskal-Wallis. **** p-value < 0.0001. a.u. = arbitrary units.

**Figure S3:**
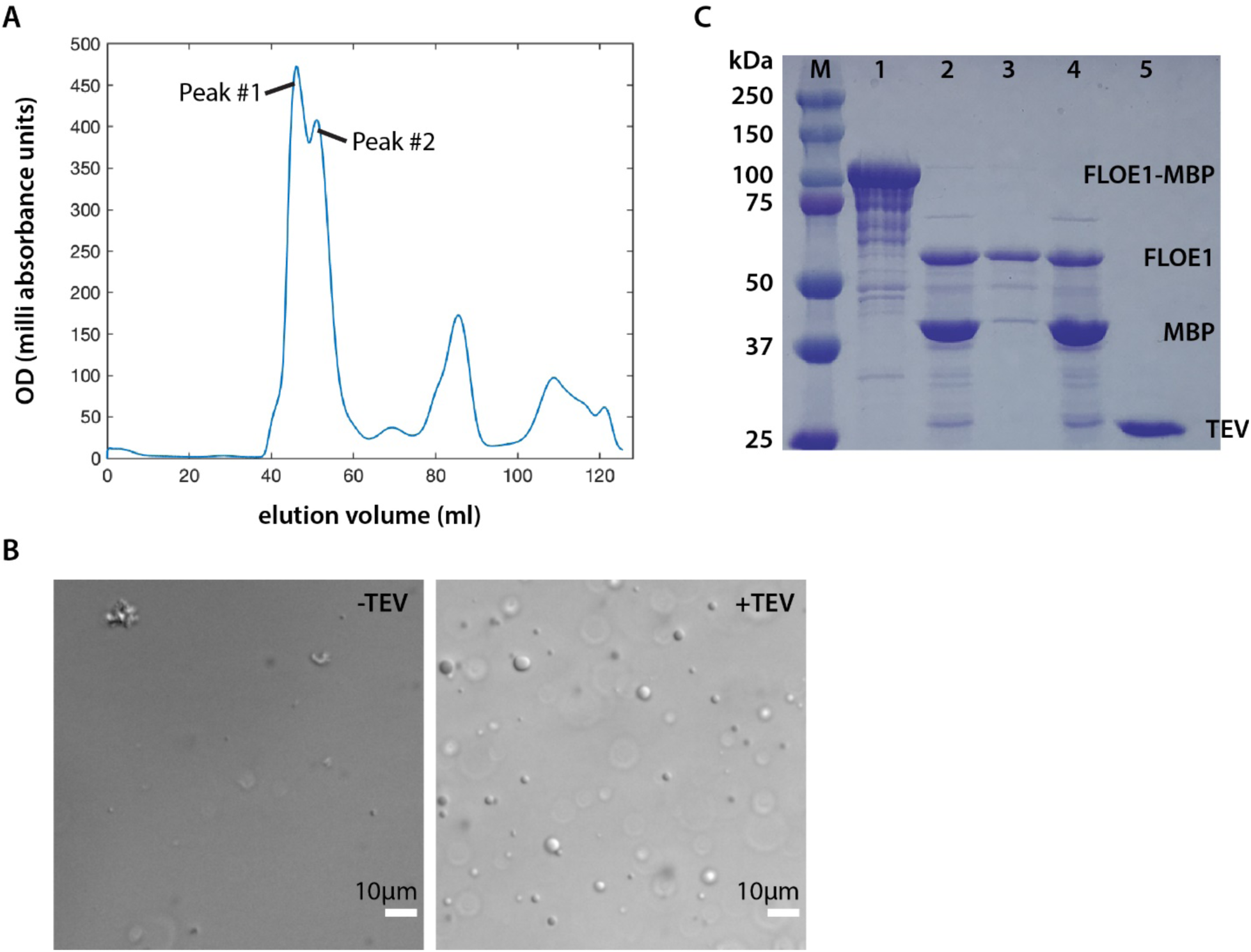
Recombinant FLOE1 expression and phase separation. (A) Size exclusion chromatography of FLOE1-MBP. For more details about Peaks # 1 and 2, see the Materials and Methods section. (B) Differential interference contrast (DIC) images of FLOE1-MBP before (left) and after (right) TEV cleavage. Some irregularly shaped aggregates can be seen in the starting material, but otherwise the sample has no droplets. After removal of the solubilizing protein, MBP, through TEV cleavage, FLOE1 forms spherical droplets indicative of liquid-liquid phase separation (LLPS). Scale bars are 10 μm. (C) SDS-PAGE confirms that TEV cleavage reaction was successful and shows the composition of the droplets observed in (B). Lane 1: FLOE1-MBP uncleaved construct (Peak #2). Lane 2: Sample from Peak #2 following 20 hours of incubation with TEV. To see the composition of the droplets and ensure they are primarily composed of FLOE1, we spun down the sample for 10 minutes under 20,000 g and separated the supernatant from the pellet. Lane 3: Pellet shows strong FLOE1 enrichment and very little MBP. Lane 4: Supernatant shows that both FLOE1 and MBP are present. The existence of a FLOE1 band in the supernatant indicates an equilibrium between the dilute and dense phases of FLOE1, as expected in LLPS. The strong MBP band indicates that these FLOE1 droplets selectively exclude MBP. Lane 5: TEV.

**Figure S4:**
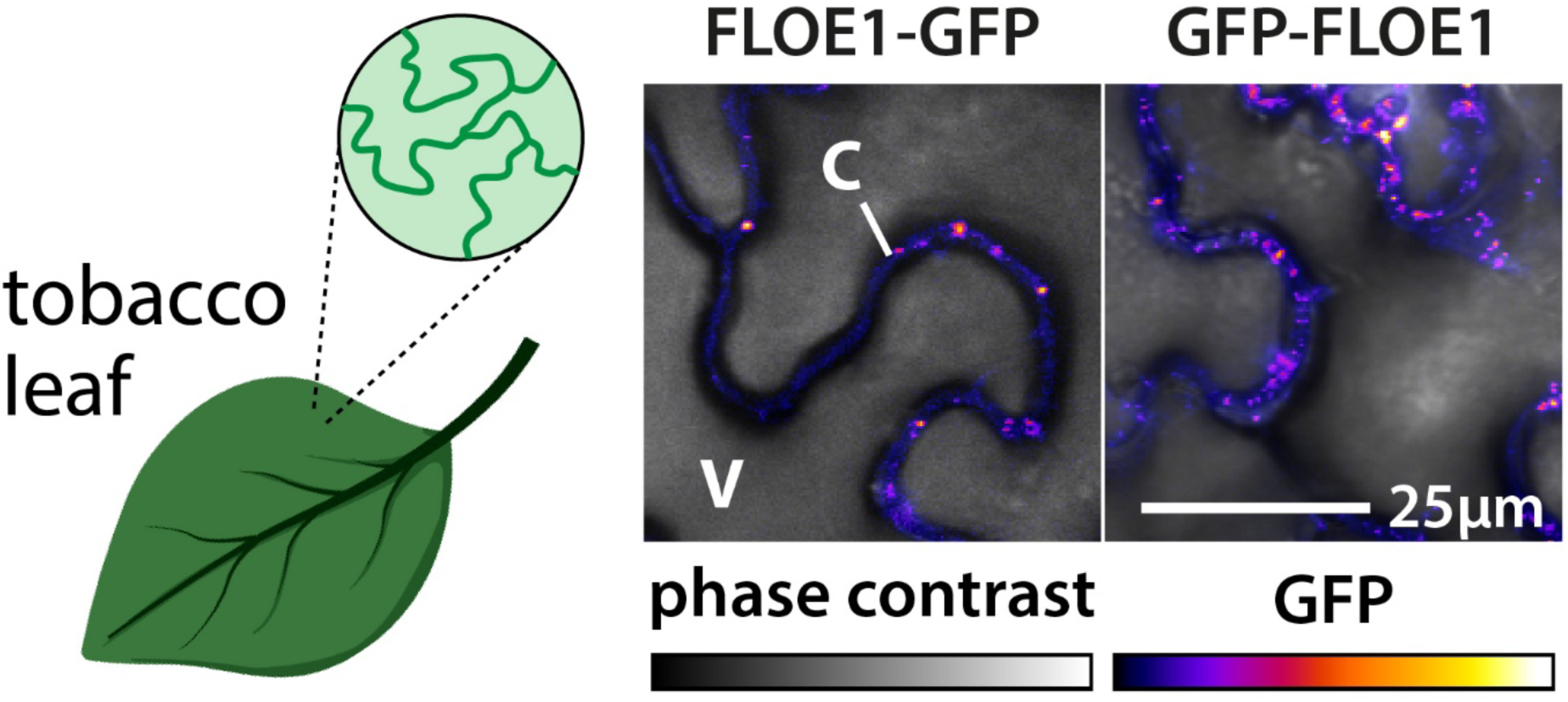
FLOE1 expression in tobacco leaves. Both N- and C-terminal GFP fusions condense into cytoplasmic condensates. V denotes vacuole, C denotes cytoplasm.

**Figure S5:**
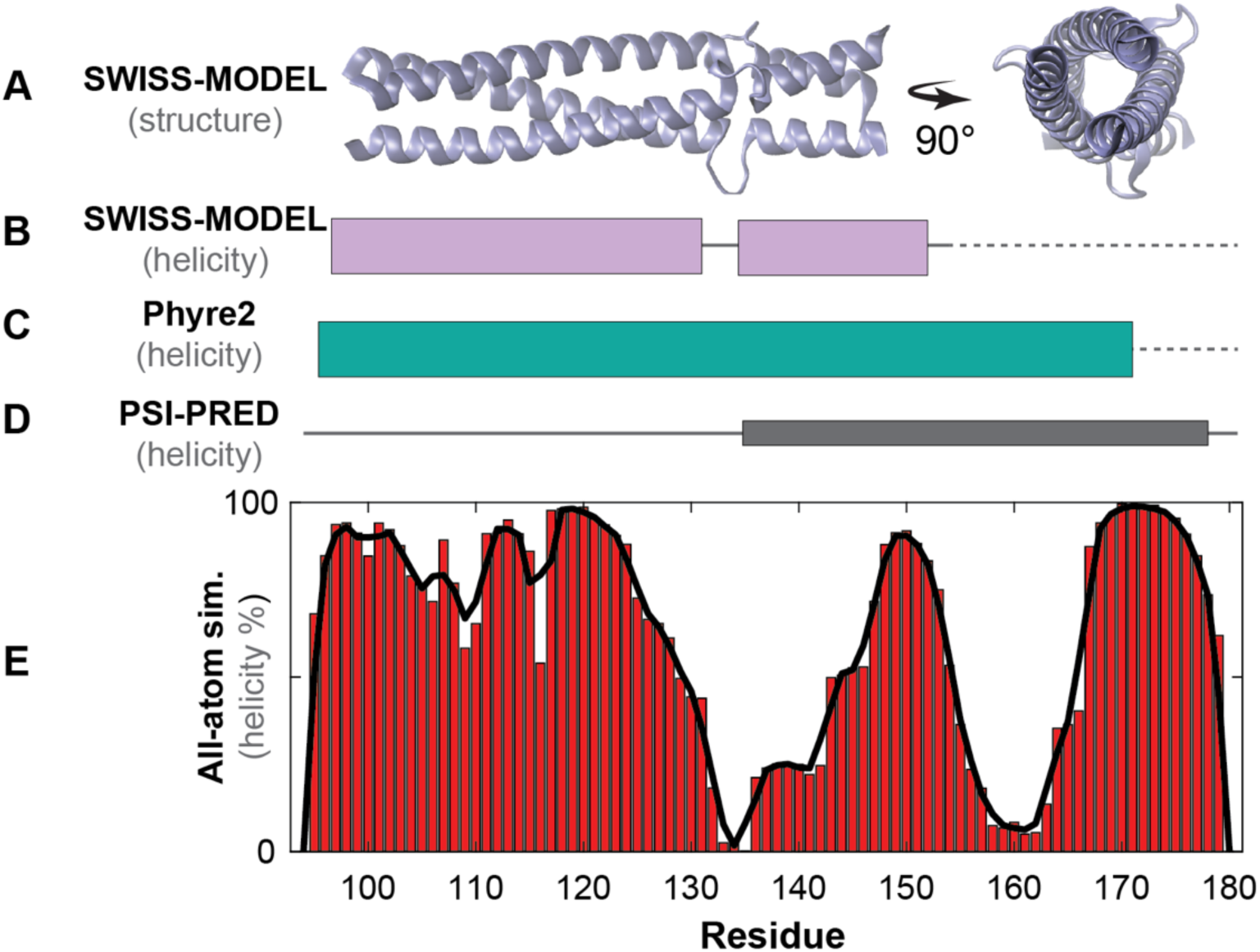
The nucleation domain is predicted to adopt a trimeric coiled-coil conformation. (A) Predicted structure from the SWISS-MODEL homology prediction server. Side and top views are shown. A well-defined trimeric coiled-coil oligomeric state is strongly predicted. (B) Inferred helicity from SWISS-MODEL prediction projected from N-to-C terminus and aligned with residue numbers in (E). (C) Predicted helicity from Phyre2 homology model prediction. (D) Predicted helicity from PSI-PRED secondary structure prediction tool. (E) Per-residue percentage helicity as obtained from all-atom simulations. Smoothened fit is shown in black.

**Figure S6:**
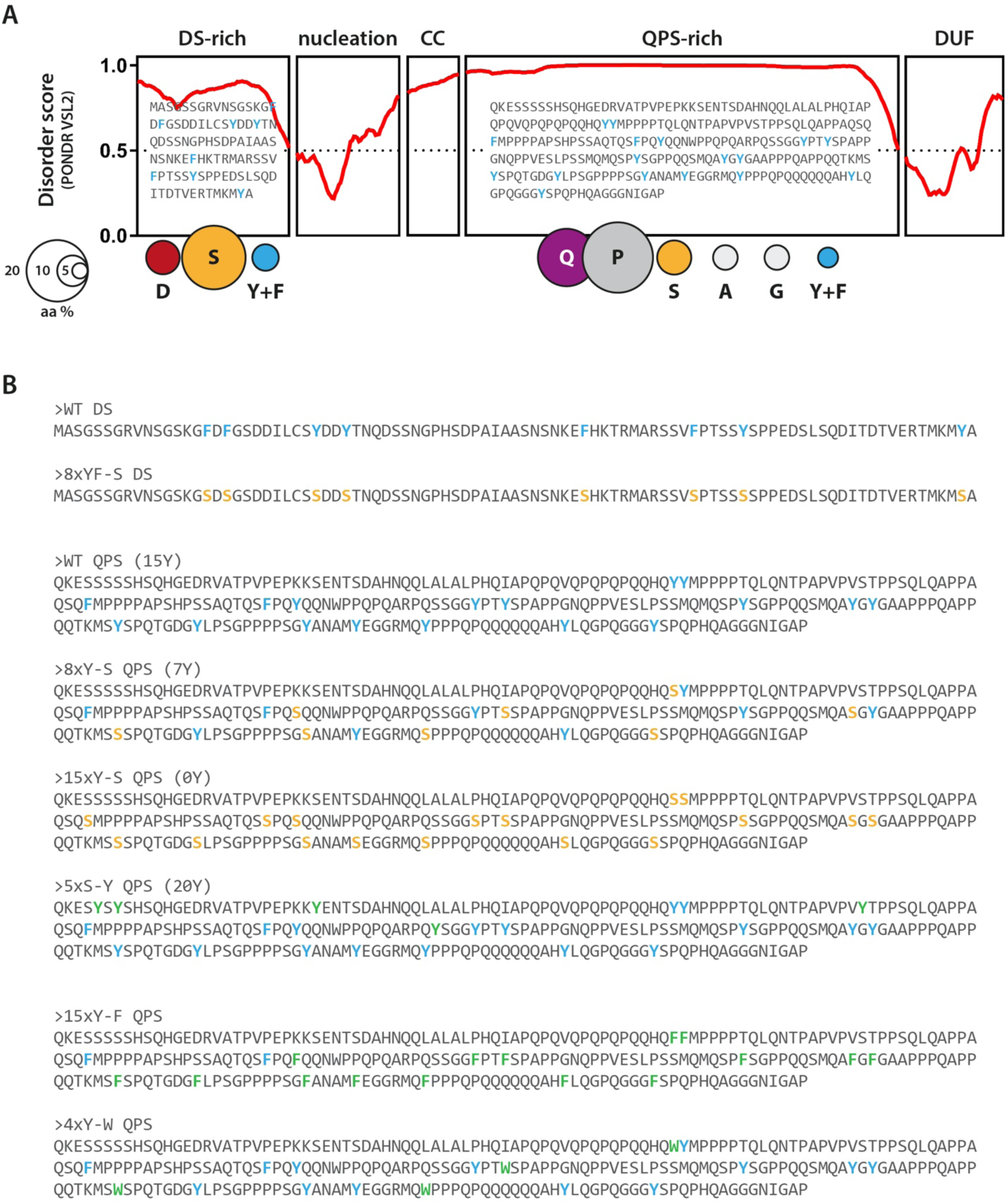
Amino acid substitution mutants. (A) Domain architecture of FLOE1 with repetitively spaced aromatic residues highlighted in blue. (B) Sequences of amino acid substitution mutants. Yellow represents serine substitutions and green represents aromatic residue substitutions. Original aromatic residues are shown in blue.

**Figure S7:**
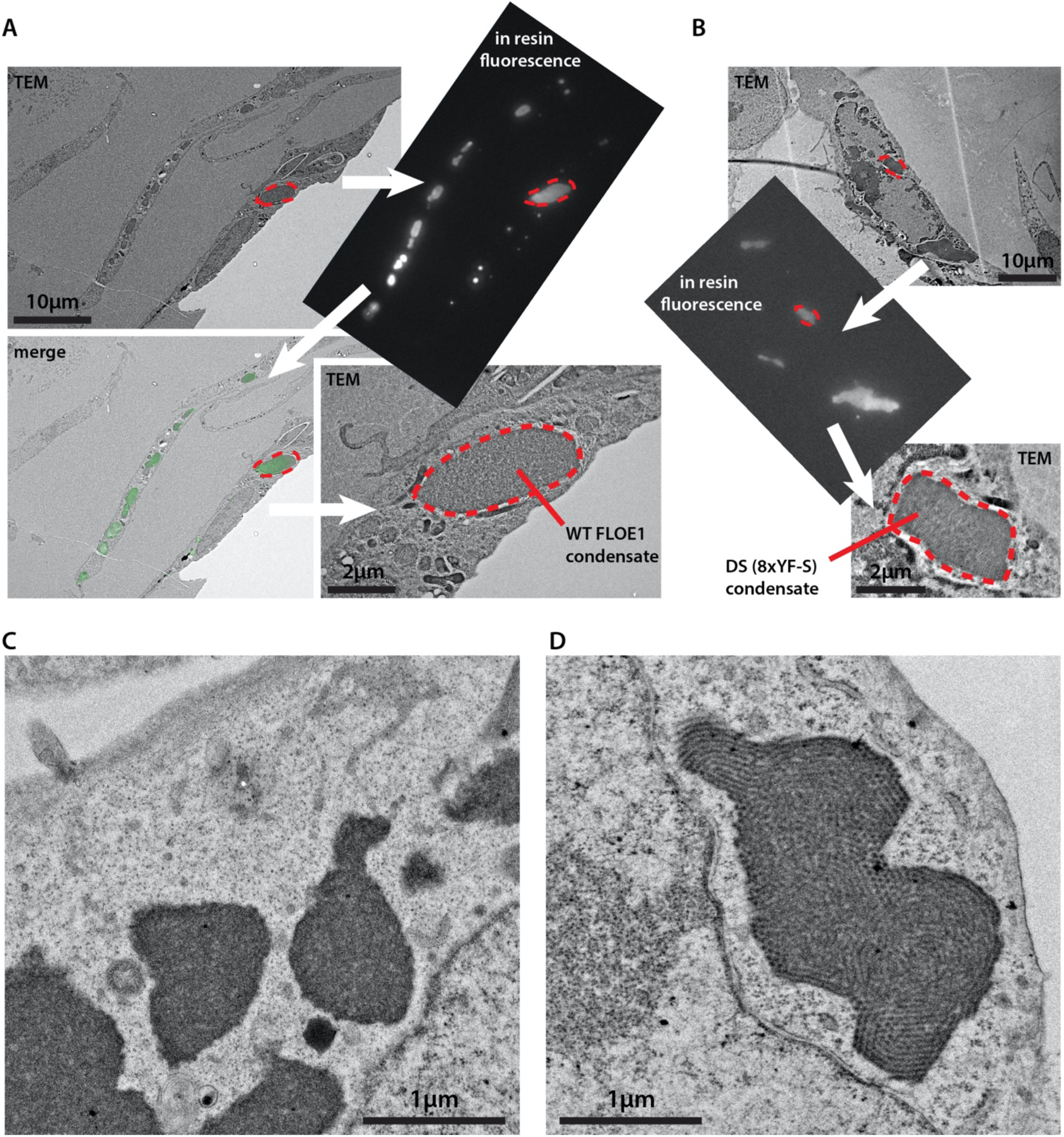
Electron microscopy to interrogate FLOE1 condensate substructure. (A-B) Correlative light-electron microscopy (CLEM) allows us to confirm the presence of GFP-FLOE1 in cytoplasmic condensates in transfected U2OS cells. (A) Shows wildtype FLOE1 condensates, whereas (B) shows mutant DS domain (8xYF-S) condensates. (C-D) Standard transmission electron microscopy of wildtype (C) and mutated DS domain (D) FLOE1 condensates. Zooms of (C-D) can be found in Fig. 2M. (D) Highlights filamentous substructure of mutated DS FLOE1 condensates.

**Figure S8:**
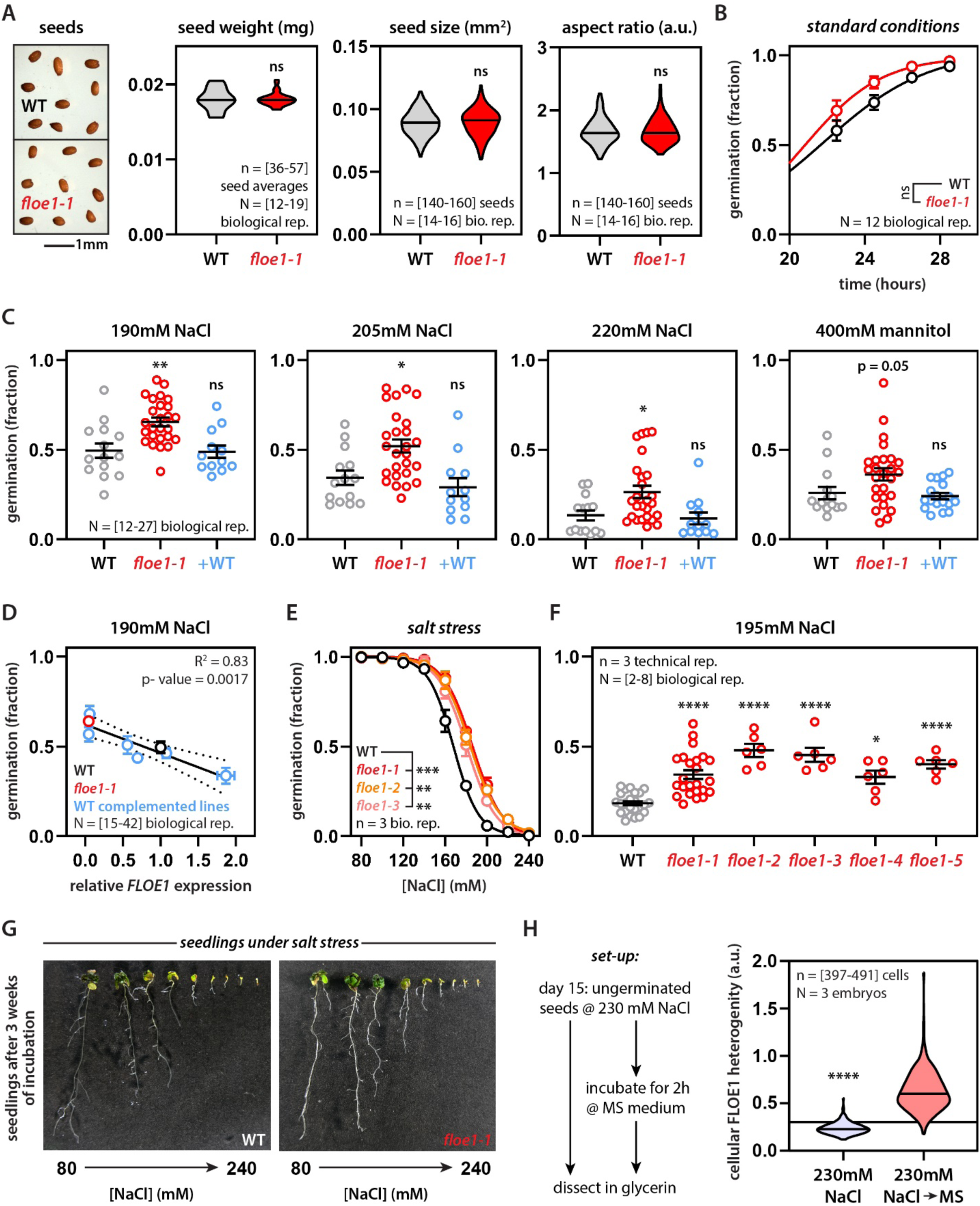
FLOE1 function modifies germination rates under water stress. (A) Absence of FLOE1 does not affect seed characteristics. Mann-Whitney. ns = not significant (B) Absence of FLOE1 does not affect germination under standard conditions. Mean ± SEM. Four-parameter dose-response fit. Two-way ANOVA with Šidák correction. Representative of two independent experiments. (C) Increased germination of *floe1-1* knockout line under water stress is rescued by WT FLOE1 complementation (+WT). Mean ± SEM. One-way ANOVA. Representative of three independent experiments. (D) Different FLOE1 WT complemented lines with different expression levels, as assayed by RT-qPCR, show dose-dependent effect of FLOE1 function on germination under salt stress. Mean ± SEM. Linear regression. (E) Two CRISPR-Cas9 *FLOE1* mutant lines show enhanced germination under varying salt stress conditions. Mean ± SEM. Four-parameter dose-response fit. Two-way ANOVA. Representative of two independent experiments. (F) Four CRISPR-Cas9 *FLOE1* mutants lines show enhanced germination under salt stress. Mean ± SEM. One-way ANOVA. Representative of three independent experiments. (G) Both WT and *floe1-1* seedlings show developmental defects upon germination under salt stress. *floe1-1* picture is the same as in Fig. 3B and is shown for comparison. (H) Quantification of FLOE1 condensate formation upon alleviation from salt stress. Horizontal line indicates cut-off for FLOE1 condensation (see Material and Methods). Mann-Whitney. * p-value < 0.05, ** p-value < 0.01, *** p-value < 0.001, **** p-value < 0.0001. MS = Murashige and Skoog medium (standard conditions). a.u. = arbitrary units.

**Figure S9:**
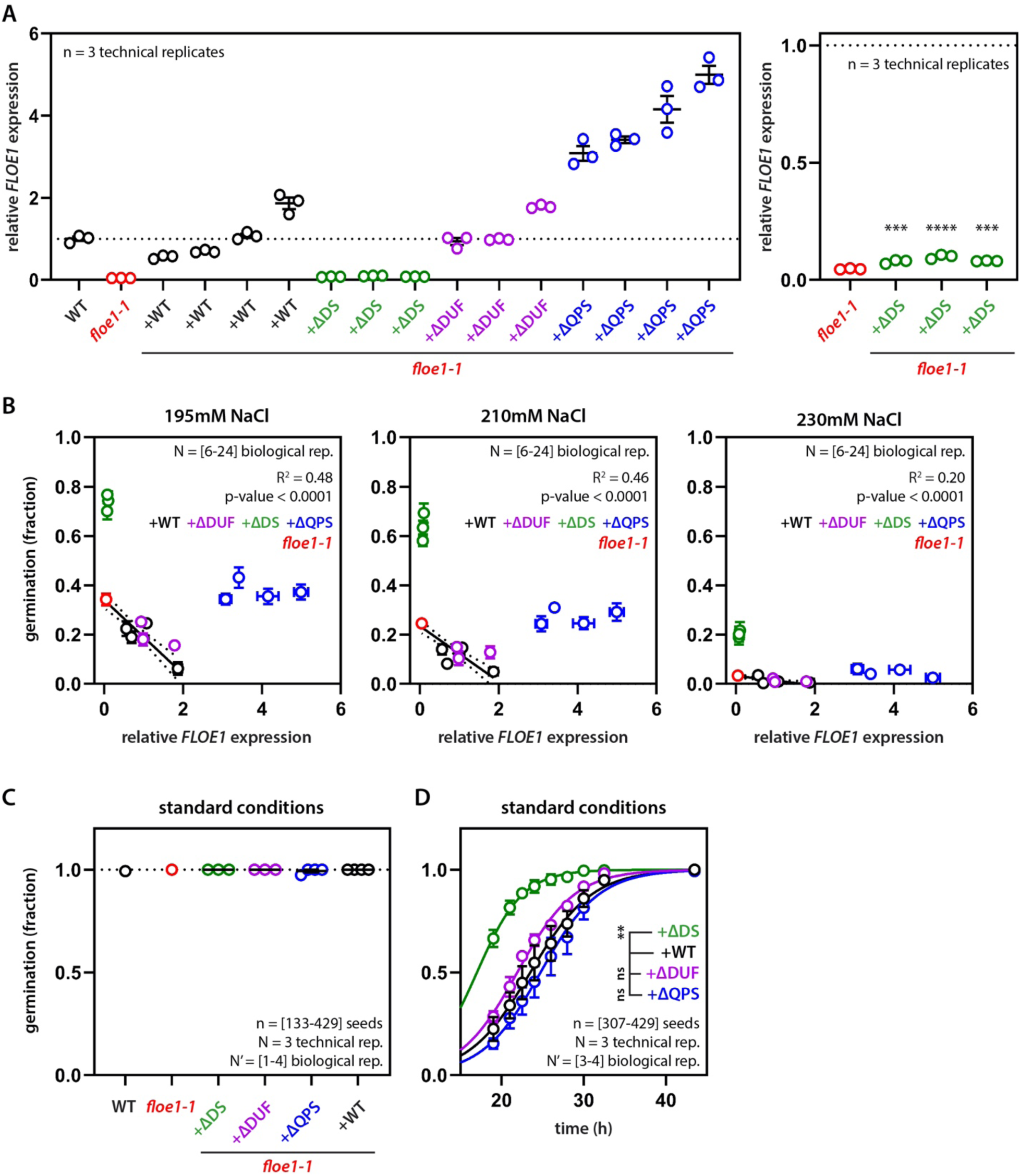
Mutant phenotypes are not due to differences in expression level. (A-B) Since FLOE1 is a dosage-dependent regulator of seed germination under water stress, we wanted to rule out that expression differences in the mutant lines would be responsible for the observed differences in their germination rates. We assayed *FLOE1* expression levels in dry seeds via RT-qPCR (A). As shown before, there was a linear correlation between *FLOE1* expression level and the germination rate (B). *floe1-1* lines complemented with the ΔDUF mutant followed a similar trend, confirming that the DUF domain deletion does not affect germination in our assays (B). *floe1-1* lines complemented with the ΔDS mutant showed low levels of transgene expression according to RT-qPCR (A, Right Panel. One-way ANOVA. *** p-value < 0.001. Mean ± SEM.) which was consistent with the sparser localization of the protein in radicles (Fig. 4F). Yet, despite these low expression levels, the ΔDS complemented lines consistently induced extreme germination rates, which we never observed for *floe1-1* or WT complemented lines. *floe1-1* lines complemented with the ΔQPS mutant showed high levels of transgene expression according to RT-qPCR (B). Despite these high transgene levels, and robust protein expression in radicles (Fig. 4F), ΔQPS complemented lines had germination rates similar to the parental *floe1-1* line, in stark contrast with WT complemented lines with higher relative expression, supporting the loss-of-function phenotype of this mutant. B: Mean ± SEM. Germination data are representative of three independent experiments. (C) All complemented lines are able to fully germinate under standard conditions (43.5h time point shown) Mean ± SEM. Representative of two independent experiments. (D) ΔDUF and ΔQPS complemented lines have similar germination rates as WT complemented lines. In contrast, ΔDS complemented lines show faster germination rates under standard conditions. Mean ± SEM. Two-way ANOVA. Average of 3-4 independent transgenic lines.

**Figure S10:**
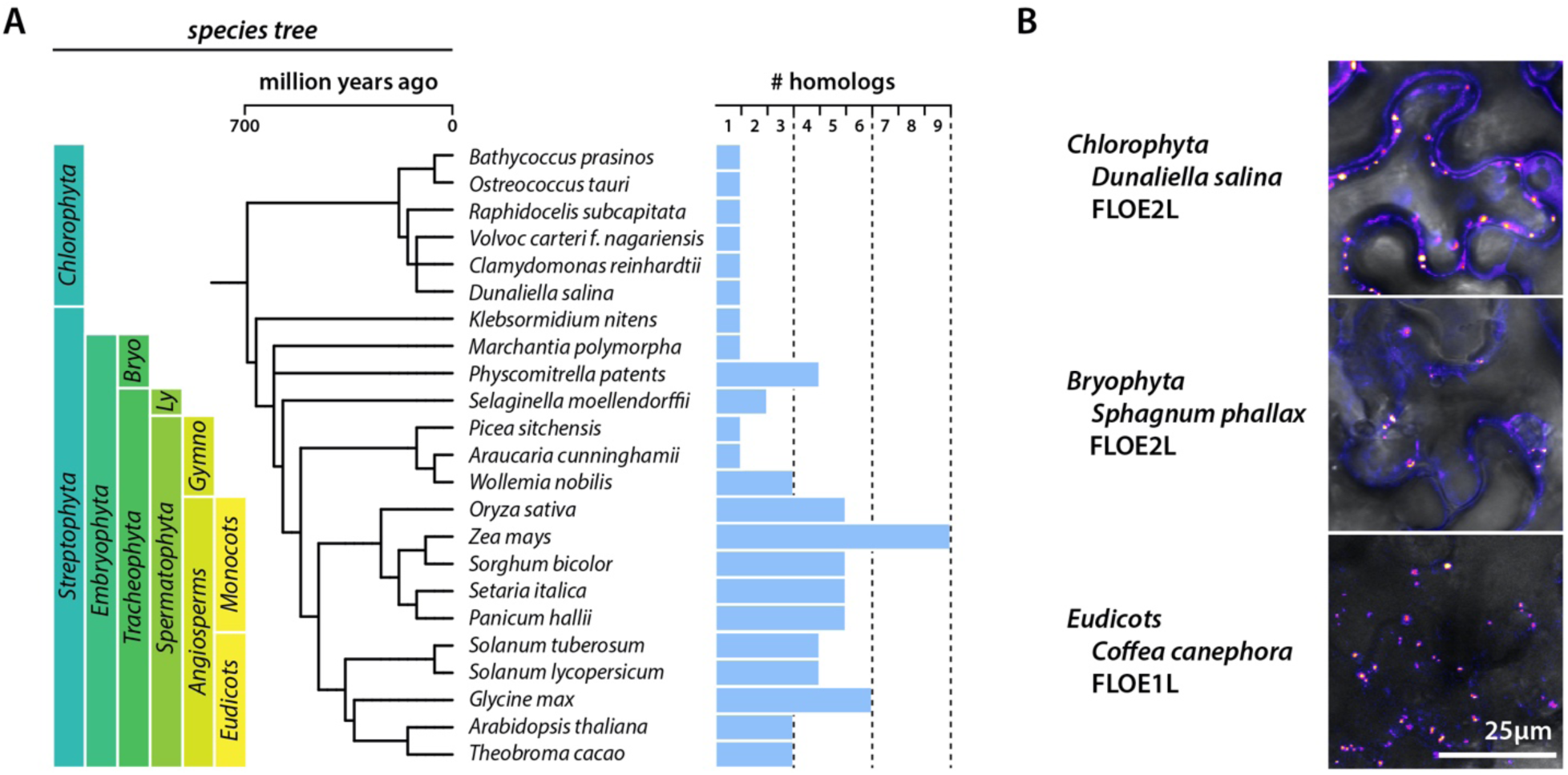
Additional information on FLOE1 homologs. (A) Species tree as in Fig. 4E with full species names. “Bryo”, “Ly” and “Gymno” refer to the bryophyte, lycophyte and gymnosperm lineages (B) Additional examples of FLOE homologs that condense upon expression in tobacco leaves. Belonging to the FLOE1L and FLOE2L clades are indicated.

**Figure S11:**
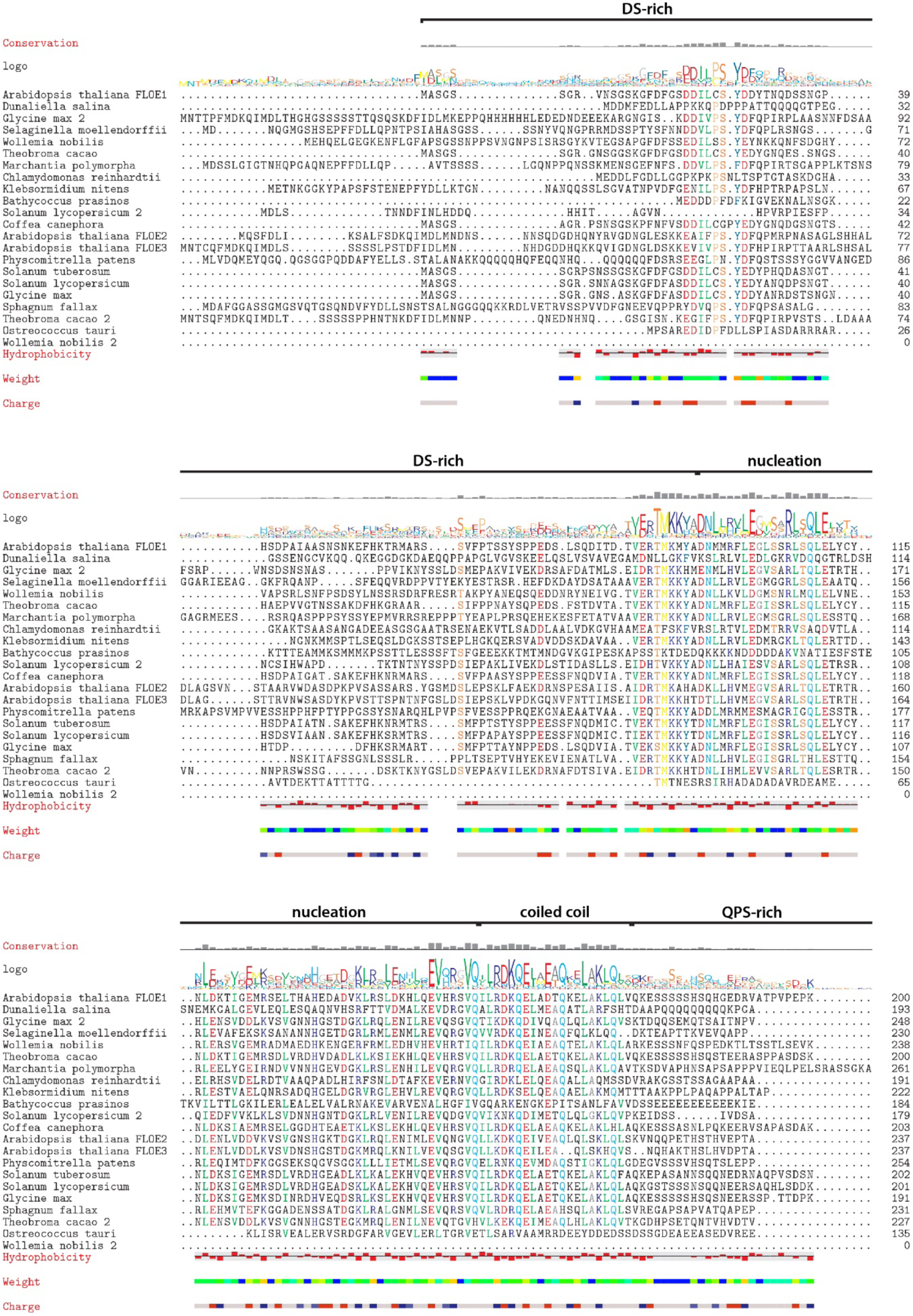
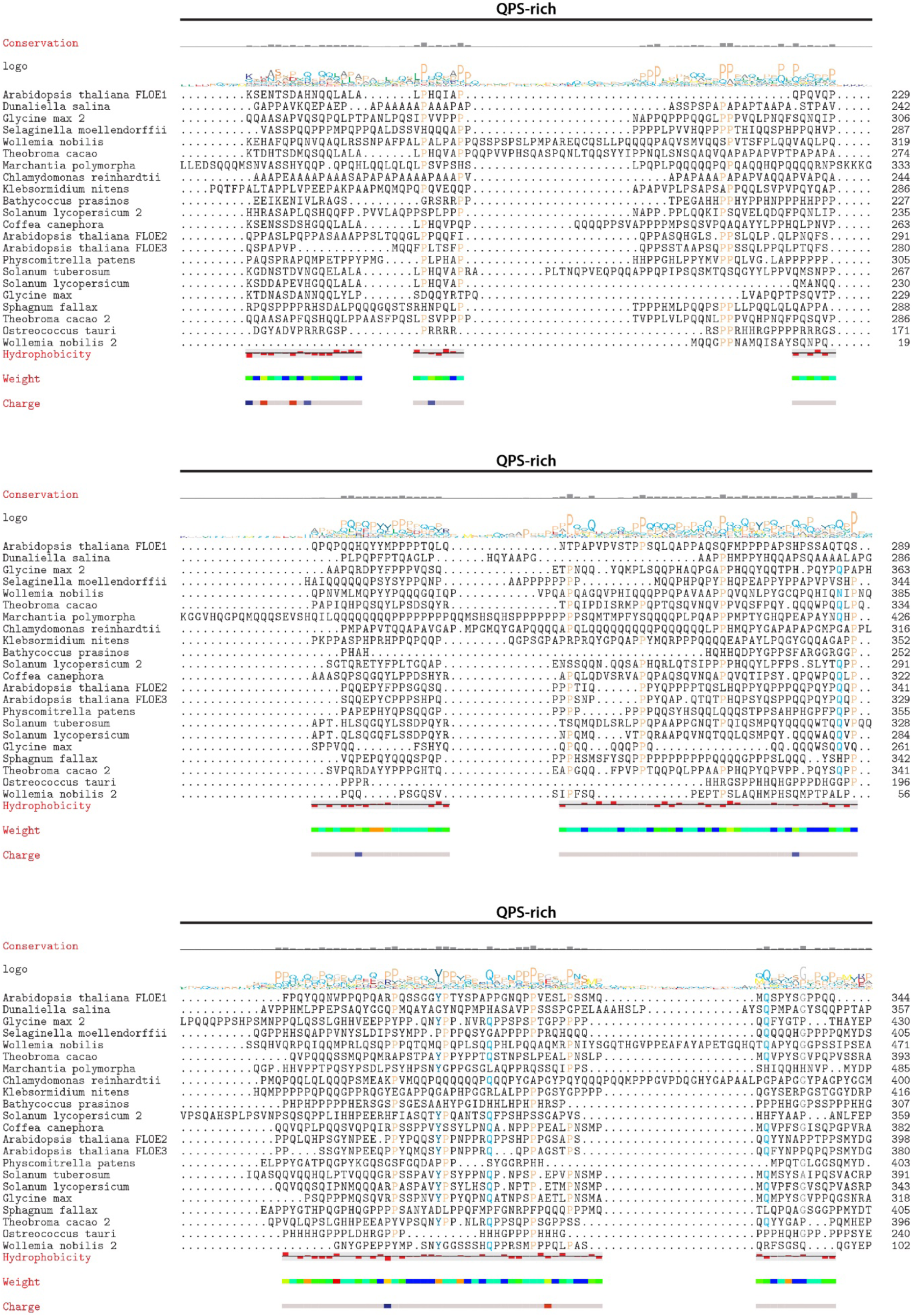
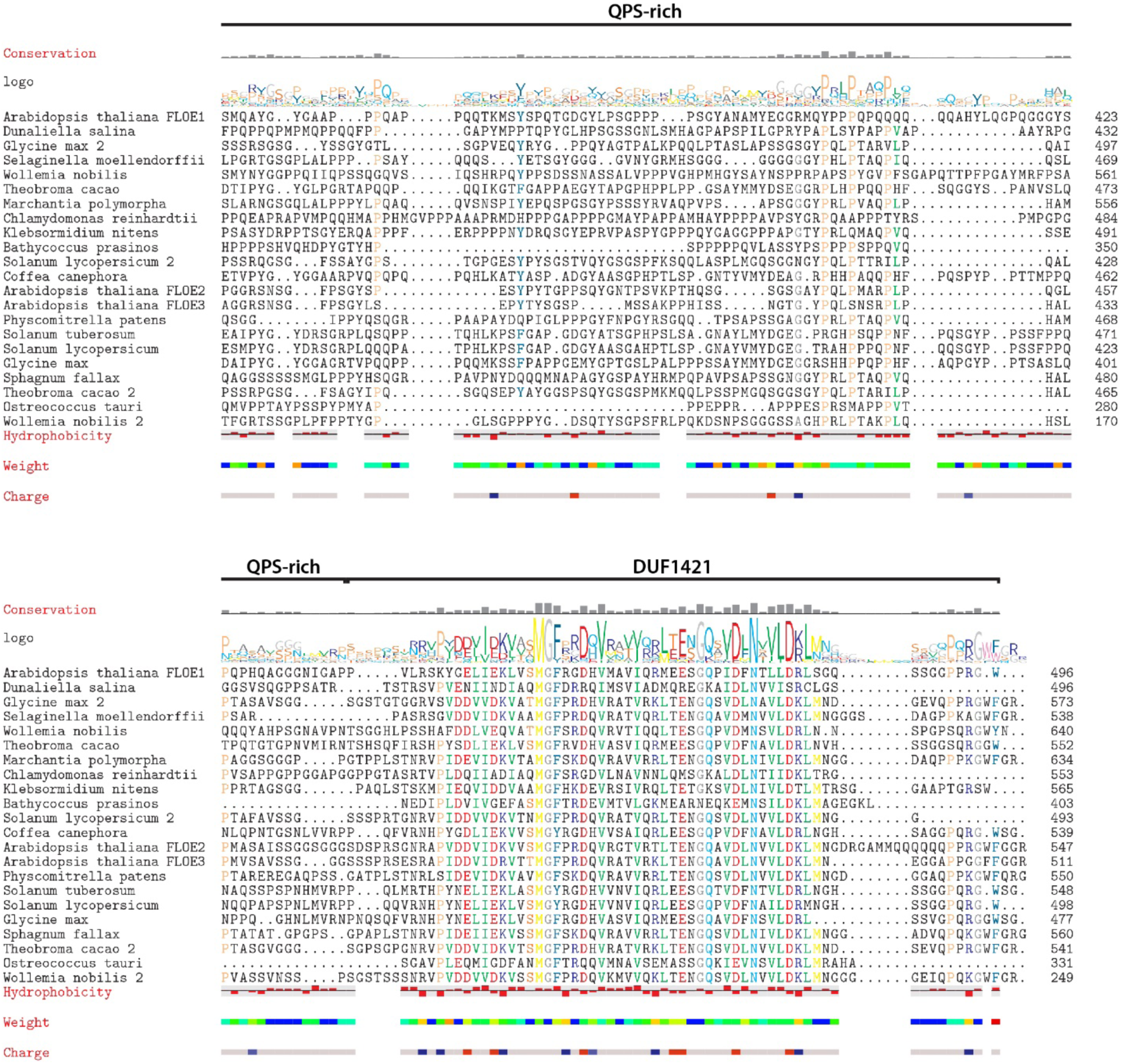
Protein sequence alignment of tested FLOE homologs. Homologs from across the plant kingdom show extensive sequence variation in both the DS and QPS disordered domains but high conservation in the other domains.

**Figure S12:**
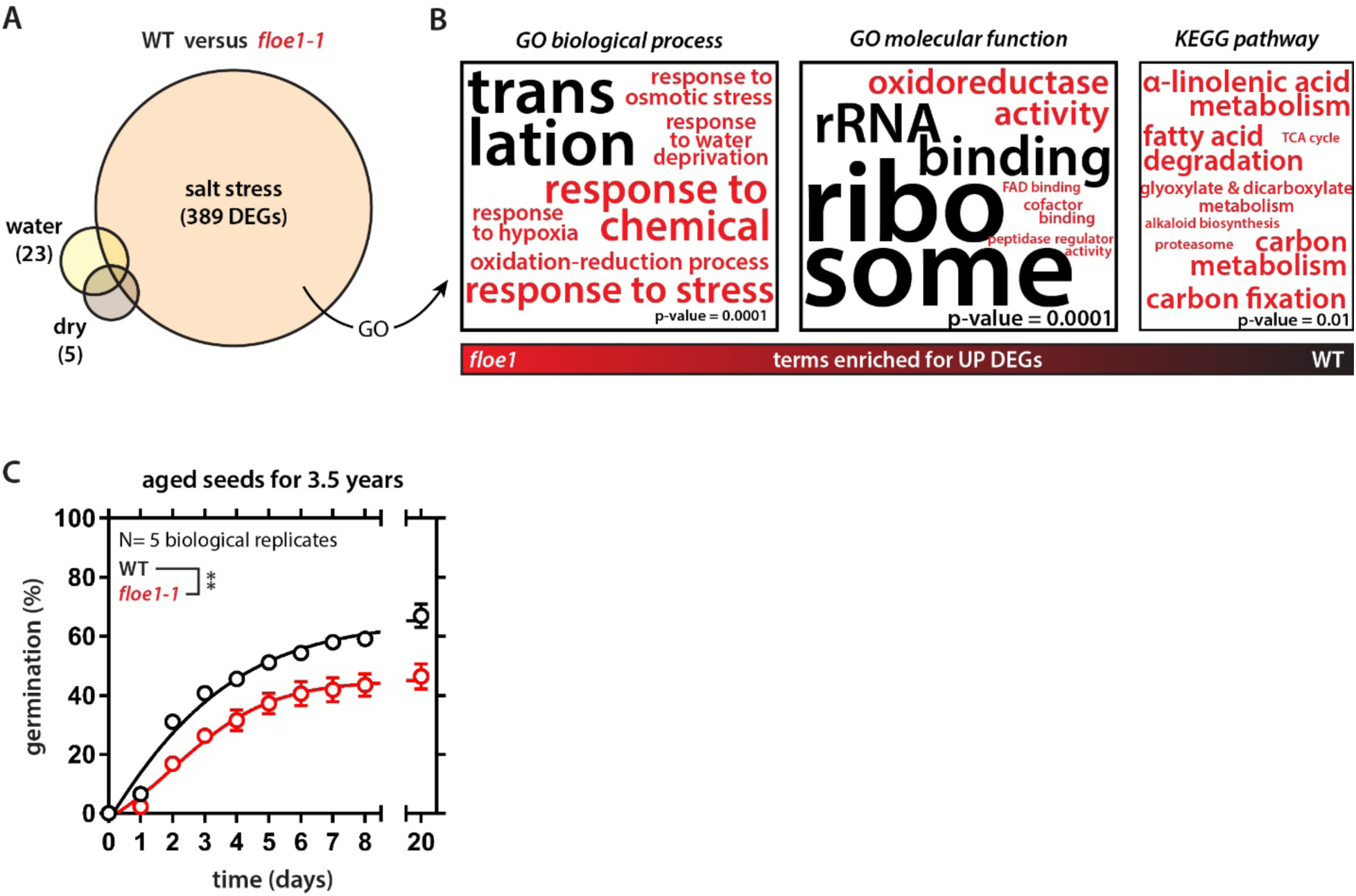
RNA seq analysis and aging of WT and *floe1-1* seeds. (A) Venn diagram showing differentially expressed genes (DEGs) between wildtype and *floe1-1* seeds under different conditions: dry seed (dry), normal imbibition (water), imbibition in 220 mM NaCl (salt stress). (B) Word cloud showing enrichment of GO or KEGG terms for DEGs under salt stress. Red terms are associated with *floe1-1* upregulated DEGs, black terms are associated with wildtype upregulated (or *floe1-1* downregulated) DEGs. Font size is proportional to –log10(p-value). The only KEGG pathway enriched for the WT was “ribosome” (p-value = 3.88E-17, not shown). (See also Table S2). (C) *floe1-1* seeds show a decreased germination potential upon aging. Average germination rates of seeds from five plants per genotype. Mean ± SEM. Four-parameter dose-response fit. Two-way ANOVA. ** p-value < 0.01.

## Notes

### Competing Interest Statement

The authors have declared no competing interest.

## REFERENCES

1. S. Yashina et al., Regeneration of whole fertile plants from 30,000-y-old fruit tissue buried in Siberian permafrost. Proc Natl Acad Sci U S A 109, 4008–4013 (2012).

2. N. Sano et al., Staying Alive: Molecular Aspects of Seed Longevity. Plant Cell Physiol 57, 660–674 (2016).

3. J. Buitink, O. Leprince, Intracellular glasses and seed survival in the dry state. C R Biol 331, 788–795 (2008).

4. O. Leprince, J. Buitink, Introduction to desiccation biology: from old borders to new frontiers. Planta 242, 369–378 (2015).

5. L. Rajjou, et al., Seed germination and vigor. Annu Rev Plant Biol 63, 507–533 (2012).

6. H. Nonogaki, F. Chen, K. J. Bradford, Mechanisms and Genes Involved in Germination Sensu Stricto. Annual Plant Reviews online 27, (2018).

7. B. Bai et al., Ecotypic variability in the metabolic response of seeds to diurnal hydration-dehydration cycles and its relationship to seed vigor. Plant Cell Physiol 53, 38–52 (2012).

8. I. Kranner, F. V. Minibayeva, R. P. Beckett, C. E. Seal, What is stress? Concepts, definitions and applications in seed science. New Phytol 188, 655–673 (2010).

9. M. Schmid et al., A gene expression map of Arabidopsis thaliana development. Nat Genet 37, 501–506 (2005).

10. R. S. Austin et al., New BAR tools for mining expression data and exploring Cis-elements in Arabidopsis thaliana. Plant J 88, 490–504 (2016).

11. Y. Shin, C. P. Brangwynne, Liquid phase condensation in cell physiology and disease. Science 357, (2017).

12. S. Boeynaems et al., Protein Phase Separation: A New Phase in Cell Biology. Trends Cell Biol 28, 420–435 (2018).

13. S. Alberti, R. Halfmann, O. King, A. Kapila, S. Lindquist, A systematic survey identifies prions and illuminates sequence features of prionogenic proteins. Cell 137, 146–158 (2009).

14. R. Halfmann et al., Prions are a common mechanism for phenotypic inheritance in wild yeasts. Nature 482, 363–368 (2012).

15. E. Gutierrez-Beltran, P. N. Moschou, A. P. Smertenko, P. V. Bozhkov, Tudor staphylococcal nuclease links formation of stress granules and processing bodies with mRNA catabolism in Arabidopsis. Plant Cell 27, 926–943 (2015).

16. M. C. Munder et al., A pH-driven transition of the cytoplasm from a fluid- to a solid-like state promotes entry into dormancy. Elife 5, (2016).

17. K. A. Burke, A. M. Janke, C. L. Rhine, N. L. Fawzi, Residue-by-Residue View of In Vitro FUS Granules that Bind the C-Terminal Domain of RNA Polymerase II. Mol Cell 60, 231–241 (2015).

18. J. Wang et al., A Molecular Grammar Governing the Driving Forces for Phase Separation of Prion-like RNA Binding Proteins. Cell 174, 688–699 e616 (2018).

19. E. W. Martin et al., Valence and patterning of aromatic residues determine the phase behavior of prion-like domains. Science 367, 694–699 (2020).

20. O. Rog, S. Kohler, A. F. Dernburg, The synaptonemal complex has liquid crystalline properties and spatially regulates meiotic recombination factors. Elife 6, (2017).

21. J. R. Gremer, D. L. Venable, Bet hedging in desert winter annual plants: optimal germination strategies in a variable environment. Ecol Lett 17, 380–387 (2014).

22. I. G. Johnston, G. W. Bassel, Identification of a bet-hedging network motif generating noise in hormone concentrations and germination propensity in Arabidopsis. J R Soc Interface 15, (2018).

23. P. Villa Martin, M. A. Munoz, S. Pigolotti, Bet-hedging strategies in expanding populations. PLoS Comput Biol 15, e1006529 (2019).

24. J. A. Riback et al., Stress-Triggered Phase Separation Is an Adaptive, Evolutionarily Tuned Response. Cell 168, 1028–1040 e1019 (2017).

25. T. M. Franzmann et al., Phase separation of a yeast prion protein promotes cellular fitness. Science 359, (2018).

26. O. D. King, A. D. Gitler, J. Shorter, The tip of the iceberg: RNA-binding proteins with prion-like domains in neurodegenerative disease. Brain Res 1462, 61–80 (2012).

27. S. K. Powers et al., Nucleo-cytoplasmic Partitioning of ARF Proteins Controls Auxin Responses in Arabidopsis thaliana. Mol Cell 76, 177–190 e175 (2019).

28. S. Chakrabortee et al., Luminidependens (LD) is an Arabidopsis protein with prion behavior. Proc Natl Acad Sci U S A 113, 6065–6070 (2016).

29. X. Fang et al., Arabidopsis FLL2 promotes liquid-liquid phase separation of polyadenylation complexes. Nature 569, 265–269 (2019).

30. B. Bakthavachalu et al., RNP-Granule Assembly via Ataxin-2 Disordered Domains Is Required for Long-Term Memory and Neurodegeneration. Neuron 98, 754–766 e754 (2018).

31. A. R. Schwember, K. J. Bradford, Quantitative trait loci associated with longevity of lettuce seeds under conventional and controlled deterioration storage conditions. J Exp Bot 61, 4423–4436 (2010).

32. T. C. Boothby, G. J. Pielak, Intrinsically Disordered Proteins and Desiccation Tolerance: Elucidating Functional and Mechanistic Underpinnings of Anhydrobiosis. Bioessays 39, (2017).

33. J. Esbelin, T. Santos, M. Hebraud, Desiccation: An environmental and food industry stress that bacteria commonly face. Food Microbiol 69, 82–88 (2018).

34. V. Giarola, Q. Hou, D. Bartels, Angiosperm Plant Desiccation Tolerance: Hints from Transcriptomics and Genome Sequencing. Trends Plant Sci 22, 705–717 (2017).

## SUPPLEMENTAL REFERENCES

1. R. S. Austin et al., New BAR tools for mining expression data and exploring Cis-elements in Arabidopsis thaliana. Plant J 88, 490–504 (2016).

2. D. Piovesan et al., MobiDB 3.0: more annotations for intrinsic disorder, conformational diversity and interactions in proteins. Nucleic Acids Res 46, D471–D476 (2018).

3. N. Xiao, D. S. Cao, M. F. Zhu, Q. S. Xu, protr/ProtrWeb: R package and web server for generating various numerical representation schemes of protein sequences. Bioinformatics 31, 1857–1859 (2015).

4. S. Chakrabortee et al., Luminidependens (LD) is an Arabidopsis protein with prion behavior. Proc Natl Acad Sci U S A 113, 6065–6070 (2016).

5. B. Xue, R. L. Dunbrack, R. W. Williams, A. K. Dunker, V. N. Uversky, PONDR-FIT: a meta-predictor of intrinsically disordered amino acids. Biochim Biophys Acta 1804, 996–1010 (2010).

6. A. K. Lancaster, A. Nutter-Upham, S. Lindquist, O. D. King, PLAAC: a web and command-line application to identify proteins with prion-like amino acid composition. Bioinformatics 30, 2501–2502 (2014).

7. S. J. Clough, Floral dip: agrobacterium-mediated germ line transformation. Methods Mol Biol 286, 91–102 (2005).

8. J. Steinert, S. Schiml, F. Fauser, H. Puchta, Highly efficient heritable plant genome engineering using Cas9 orthologues from Streptococcus thermophilus and Staphylococcus aureus. Plant J 84, 1295–1305 (2015).

9. T. Nakagawa et al., Development of series of gateway binary vectors, pGWBs, for realizing efficient construction of fusion genes for plant transformation. J Biosci Bioeng 104, 34–41 (2007).

10. R. V. Davuluri et al., AGRIS: Arabidopsis gene regulatory information server, an information resource of Arabidopsis cis-regulatory elements and transcription factors. BMC Bioinformatics 4, 25 (2003).

11. F. Bossi et al., Systematic discovery of novel eukaryotic transcriptional regulators using sequence homology independent prediction. BMC Genomics 18, 480 (2017).

12. C. UniProt, UniProt: a worldwide hub of protein knowledge. Nucleic Acids Res 47, D506–D515 (2019).

13. D. M. Goodstein et al., Phytozome: a comparative platform for green plant genomics. Nucleic Acids Res 40, D1178–1186 (2012).

14. F. Lemoine et al., NGPhylogeny.fr: new generation phylogenetic services for non-specialists. Nucleic Acids Res 47, W260–W265 (2019).

15. I. Letunic, P. Bork, Interactive Tree Of Life (iTOL) v4: recent updates and new developments. Nucleic Acids Res 47, W256–W259 (2019).

16. U. Bodenhofer, E. Bonatesta, C. Horejs-Kainrath, S. Hochreiter, msa: an R package for multiple sequence alignment. Bioinformatics 31, 3997–3999 (2015).

17. E. Beitz, TEXshade: shading and labeling of multiple sequence alignments using LATEX2 epsilon. Bioinformatics 16, 135–139 (2000).

18. R. A. Jefferson, T. A. Kavanagh, M. W. Bevan, GUS fusions: beta-glucuronidase as a sensitive and versatile gene fusion marker in higher plants. EMBO J 6, 3901–3907 (1987).

19. J. Schindelin et al., Fiji: an open-source platform for biological-image analysis. Nat Methods 9, 676–682 (2012).

20. C. A. Schneider, W. S. Rasband, K. W. Eliceiri, NIH Image to ImageJ: 25 years of image analysis. Nat Methods 9, 671–675 (2012).

21. L. Meng, L. Feldman, A rapid TRIzol-based two-step method for DNA-free RNA extraction from Arabidopsis siliques and dry seeds. Biotechnol J 5, 183–186 (2010).

22. C. Mizzotti et al., Time-Course Transcriptome Analysis of Arabidopsis Siliques Discloses Genes Essential for Fruit Development and Maturation. Plant Physiol 178, 1249–1268 (2018).

23. B. J. Dekkers et al., Identification of reference genes for RT-qPCR expression analysis in Arabidopsis and tomato seeds. Plant Cell Physiol 53, 28–37 (2012).

24. T. Czechowski, M. Stitt, T. Altmann, M. K. Udvardi, W. R. Scheible, Genome-wide identification and testing of superior reference genes for transcript normalization in Arabidopsis. Plant Physiol 139, 5–17 (2005).

25. E. Afgan et al., The Galaxy platform for accessible, reproducible and collaborative biomedical analyses: 2018 update. Nucleic Acids Res 46, W537–W544 (2018).

26. M. I. Love, W. Huber, S. Anders, Moderated estimation of fold change and dispersion for RNA-seq data with DESeq2. Genome Biol 15, 550 (2014).

27. U. Raudvere et al., g:Profiler: a web server for functional enrichment analysis and conversions of gene lists (2019 update). Nucleic Acids Res 47, W191–W198 (2019).

28. A. Waterhouse et al., SWISS-MODEL: homology modelling of protein structures and complexes. Nucleic Acids Res 46, W296–W303 (2018).

29. L. A. Kelley, S. Mezulis, C. M. Yates, M. N. Wass, M. J. Sternberg, The Phyre2 web portal for protein modeling, prediction and analysis. Nat Protoc 10, 845–858 (2015).

30. L. J. McGuffin, K. Bryson, D. T. Jones, The PSIPRED protein structure prediction server. Bioinformatics 16, 404–405 (2000).

31. A. Vitalis, R. V. Pappu, ABSINTH: a new continuum solvation model for simulations of polypeptides in aqueous solutions. J Comput Chem 30, 673–699 (2009).

32. S. Nallamsetty, B. P. Austin, K. J. Penrose, D. S. Waugh, Gateway vectors for the production of combinatorially-tagged His6-MBP fusion proteins in the cytoplasm and periplasm of Escherichia coli. Protein Sci 14, 2964–2971 (2005).

33. K. A. Burke, A. M. Janke, C. L. Rhine, N. L. Fawzi, Residue-by-Residue View of In Vitro FUS Granules that Bind the C-Terminal Domain of RNA Polymerase II. Mol Cell 60, 231–241 (2015).

34. S. Boeynaems, M. De Decker, P. Tompa, L. Van Den Bosch, Arginine-rich Peptides Can Actively Mediate Liquid-liquid Phase Separation. Bio-protocol 7, doi: 10.21769/BioProtoc.2525 (2017).

35. K. L. McDonald, R. I. Webb, Freeze substitution in 3 hours or less. J Microsc 243, 227–233 (2011).

36. C. J. Peddie et al., Correlative and integrated light and electron microscopy of in-resin GFP fluorescence, used to localise diacylglycerol in mammalian cells. Ultramicroscopy 143, 3–14 (2014).

37. J. Waese et al., ePlant: Visualizing and Exploring Multiple Levels of Data for Hypothesis Generation in Plant Biology. Plant Cell 29, 1806–1821 (2017).

